# Identification of Functional Genetic Components Modulating Toxicity Response to PFOS using Genome-wide CRISPR Screens in HepG2/C3A cells

**DOI:** 10.64898/2025.12.16.694451

**Authors:** Chanhee Kim, Abderrahmane Tagmount, Zhaohan Zhu, Frances Wilson, Danmeng Li, David A. Ostrov, W. Brad Barbazuk, Rhonda Bacher, Chris D. Vulpe

## Abstract

Perfluorooctane sulfonate (PFOS) poses significant health and environmental risks due to its persistence and widespread use and has been linked to various adverse outcomes, such as liver toxicity. Although the molecular responses and toxicity effects of PFOS exposure have been extensively studied, considerable uncertainty remains regarding the causal mechanisms leading to PFOS-associated adverse effects. To help bridge this gap, we conducted CRISPR screens in HepG2/C3A human liver cells exposed to IC_25_ (170 µM) of PFOS to identify genes and pathways influencing PFOS-induced cytotoxicity. Using a genome-wide CRISPR knockout library targeting 18,819 genes, we identified 340 candidate genes that modulate PFOS-induced cytotoxicity when genetically disrupted (189 gene disruptions increased sensitivity and 151 gene disruptions increased resistance). From these candidate genes, we individually disrupted two candidate genes, *SLC6A9* which encodes the glycine transporter GlyT1, and *CPSF2*, and confirmed increased resistance to PFOS exposure. Further, molecular docking analysis predicts that PFOS directly binds to GlyT1 and functional inhibition of GlyT1 also increases resistance to PFOS exposure. Gene-Disease outcome association analysis using the Comparative Toxicogenomics Database (CTD) indicated an enrichment of candidate genes associated with cancer-related and liver disease phenotypes. KEGG and STRING enrichment analyses found over representation of several biological pathways including DNA damage response and cell cycle. Lastly, cross-species conservation analysis using the top two validated gene targets found that their pathways were highly conserved in several environmentally relevant species. These findings provide new mechanistic and functional insights into PFOS-induced cytotoxicity, highlight potential molecular targets for toxicity mitigation, and establish a foundation for cross-species toxicogenomic modeling of PFOS health effects.

## 1. Introduction

The number of chemicals used in consumer and industrial products has increased at a pace faster than our ability to assess the potential risks they may pose to human and ecological health. Among these, a large group known as poly- and perfluoroalkyl substances (PFAS) has raised significant concerns due to widespread use and their resistance to degradation, making them ubiquitous in the environment. They have been linked to adverse human health and environmental effects(Wee and Aris, 2023). Although its manufacture has been restricted across the globe, perfluorooctane sulfonate (PFOS), remains the most commonly detected PFAS globally and exposure represents an ongoing concern in human and ecological health(Göckener *et al*., 2020; Cheng *et al*., 2022; Wee and Aris, 2023) as evidenced by a recent large-scale environmental survey done by US EPA (Environmental Protection Agency) detecting PFOS at levels higher than US guidelines in 31 % of global groundwater samples(‘pfos-report-2024.pdf’, no date).

Several human epidemiological studies have indicated potential links between exposure to PFOS and human adverse outcomes, including reproductive and developmental defects, immunotoxicity, and, the focus of this study, liver toxicity(Cui *et al*., 2009; Salihovic *et al*., 2019; Fenton *et al*., 2021; India-Aldana *et al*., 2023). Previous research examining PFOS accumulation in human tissues revealed its presence in the brain, kidneys, liver, and lungs, with the highest concentration found in the liver(Pérez *et al*., 2013; Zeng *et al*., 2019). A systematic review and meta-analysis of human epidemiology studies found correlation between predicted or measured PFOS exposure and higher serum alanine aminotransferase (ALT) levels, a biomarker of liver injury(Borghese *et al*., 2022; Costello *et al*., 2022). Although most of the epidemiological studies suggest a potential association between PFOS exposure and liver toxicity, a causal link is still controversial. In rodent studies, PFOS treatment altered lipid utilization pathways, increasing hepatic lipid accumulation, in turn, hepatic steatosis(Marques *et al*., 2020; Pfohl *et al*., 2021; Ling *et al*., 2023). In addition, the combination of transcriptomics and proteomics revealed enrichment of hepatic steatosis-related pathways in mice exposed to PFOS in their diet, consistent with PFOS-induced liver damage(Zhang *et al*., 2023). An *in vitro* study that measured cellular triglyceride and cholesterol levels in human HepaRG cells exposed to PFOS supported PFOS-induced liver damage(Louisse *et al*., 2020). While current evidence suggests that liver toxicity can result from PFOS exposure, the molecular mechanisms and specific targets are not clearly understood.

Omics-based approaches have identified cellular pathways and putative molecular targets underlying PFOS effects(Rowan-Carroll *et al*., 2021). While these ’omics’ analyses do identify molecular changes correlated with PFOS exposure, they do not provide direct causal evidence for specific targets and/or molecular mechanisms underlying cellular- and organismal-effects of PFOS. Genome-wide CRISPR screens can provide a complementary approach to help identify the key functional genetic components and pathways modulating PFAS toxicity. In this study, we carried out a genome-wide loss-of-function CRISPR screen in human HepG2/C3A liver cells to identify functional modulators of PFOS cellular toxicity. We independently validated the effect of targeted disruption of two selected top candidate genes. In addition, an inhibitor study and molecular docking prediction were performed with one of the validated genes based on structural similarity of PFOS and an inhibitor of the gene product(de Carvalho *et al*., 2024), which suggested mechanistic insights into PFOS-induced cellular toxicity. The Comparative Toxicogenomics Database (CTD) was used to identify potential linkages to adverse outcomes (diseases) associated with the candidate genes identified in our screens. KEGG and STRING enrichment analyses of candidate genes that increase or decrease sensitivity to PFOS-induced cytotoxicity, when genetically disrupted, identified cellular processes and pathways, consistent with gene-disease association identified in CTD analysis. The top two candidate genes that were experimentally validated were further analyzed for their evolutionary conservation and functional relevance across species to provide insights into shared mechanisms underlying the adverse environmental and ecological impacts of PFOS exposure, which could guide future one-health perspective studies.

## 2. Material and methods

### 2.1. Cell cultures and PFOS cytotoxicity

HEK293T and HepG2/C3A cells were purchased from the American Type Culture Collection (ATCC, Manassas, Virginia). Cells were maintained and passaged following ATCC’s recommended protocol (for the detailed condition, see Supplementary Information). Cytostasis/Cytotoxicity of PFOS exposure in HepG2/C3A cells exposed to a range of nominal concentrations (0–300 µM) in triplicate for 6 days was assessed by measuring ATP levels using the CellTiter-Glo2.0 cell viability assay kit (Promega, Madison, WI) following the manufacturer’s instruction. Of note, we chose the duration of exposure as 6 days for the IC_25_ calculation since PFOS exposure didn’t show detectable cytotoxicity in HepG2/C3A cells after 3 days of exposure, which is the exposure period that we routinely use to estimate sublethal concentrations for CRISPR screens. Additionally, we performed a pre-screen time-course treatment with the initial benchmark IC_25_ (6 days) dose over 18 days by passaging the treated cells every 6 days to refine the exposure conditions for the screen and to assess longer-term cellular responses (including cumulative cytostasis and mild cytotoxicity) by decreasing IC_25_ (6 days) down to 170 µM as higher cytotoxicity was observed with IC_25_ (6 days) for 18 days.

### 2.2. Genome-wide CRISPR loss-of-function screening

For the genome-wide loss-of-function genetic screens, we used a customized minimal genome-wide sgRNA library (Table S1) termed MinLibCas9 based on a published study which selected two sgRNAs with maximal on-target activity for each gene based on computational analysis of the screening results of several genome-wide screens(Gonçalves *et al*., 2021). Lentivirus production was carried out as previously described(Sobh *et al*., 2019). A pool of the The MinLibCas9 lentiviral library was transduced into HepG2/C3A cells at a0.3 multiplicity of infection (MOI) to generate a mutant pool as in previous studies(Russo *et al*., 2020; Zhao *et al*., 2021) (detailed methods available in Supplementary Information). The mutant pool was then expanded for the PFOS exposure experiments to obtain sufficient number of cells for replicates and controls. Control and PFOS exposure samples of 16 X 10^6^ cells in T225 flasks (three replicates) were used for screening. Exposure samples were treated with PFOS at inhibitory concentration (IC_25_) for 18 days (10 doubling times). Representation of the library was maintained throughout the screening experiment at approximately 400X the MinLibCas9 library size (approximately 40,000, so 400X = 16,000,000 cells) for each replicate.

### 2.3. DNA extraction, library preparation, and next-generation sequencing (NGS)

Genomic DNA was extracted from 16 X 10^6^ cells of each sample using the Quick-DNA Midiprep Plus Kit (ZYMO Research) following the manufacturer’s protocol. Amplicons for NGS Illumina sequencing were generated using the pairs of universal CRISPR-FOR1 forward primer and CRISPR-REV**#** reverse primers (**#**: 1 to 48) specific for each sample (see Table S2). Amplicons amplifying the sgRNA region in each sample were then pooled and gel purified using the QIAquick Gel Extraction Kit (Qiagen) and quantified using the Qubit HS dsDNA assay (Thermo Scientific). Equimolar amounts of each amplicon library were multiplexed in one pool. The NGS was carried out at the Interdisciplinary Center for Biotechnology Research (ICBR), University of Florida at Gainesville, using the NovaSeqX paired 150 bp high-throughput platform (Illumina).

### 2.4. Data processing and bioinformatics

After data acquisition (raw fastq.gz files), read quality was checked using FASTQC tools. A Model-based Analysis of Genome-wide CRISPR-Cas9 Knockout (MaGeCK)(Li *et al*., 2014) was then used to demultiplex raw fastq data files, which were further processed to generate reads containing only the unique 20 bp guide sequences. The resulting read counts from different samples of control and PFOS exposure were then normalized to adjust for the effect of library sizes and read count distributions. The sgRNAs were then ranked by a modified robust ranking aggregation (α-RRA) algorithm, identifying candidate genes. Candidate genes were defined as genes with *p*-value < 0.01 and Log_2_FC < - 0.6 (sensitive) or Log_2_FC > 0.6 (resistant) for further analysis. We define sensitive genes as those that when disrupted by CRISPR targeting, show increased cellular sensitivity (less abundant than control) to PFOS exposure whereas resistant genes show increased resistance (more abundant than control) to PFOS exposure. We compared the MaGeCK-produced results with the results obtained using Platform-independent Analysis of Pooled Screens using Python (PinAPL-Py)(Spahn *et al*., 2017).

### 2.5. Single gene knockout validations

We performed individual disruptions of the top two candidate genes that increase resistance to PFOS when genetically disrupted. The sgRNAs targeting the *SLC6A9* and *CPSF2* genes were selected from the MinLibCas9 sgRNA library (Table S3). Each sgRNA was cloned into the LentiCRISPRv2 using the golden gate assembly strategy as described elsewhere in detail(Joung *et al*., 2017). After cloning of each sgRNA, production and transduction of the corresponding lentivirus were conducted, generating knockout cell pools (for details, see Supplementary Information). The cellular pools for each targeted gene were used for cytotoxicity assays to evaluate their resistance to PFOS exposure.

### 2.6. Molecular docking

A Protein Data Bank (PDB) format file for the GlyT1 and glycine transporter 2 (GlyT2) structures were obtained from AlphaFold Protein Structure Databse(Jumper *et al*., 2021; Varadi *et al*., 2024). Molecular docking to predict the binding of PFOS to these transporters was performed using AutoDock Vina (<u>AutoDock Vina</u>). The docking search grid was defined by a box measuring 84 × 48 × 78 Å. The highest scoring poses, indicated by the lowest ΔG values (in kcal/mol), were identified in the central region of each transporter. All structural figures were created using PyMOL (The PyMOL Molecular Graphics System, Version 3.0, Schrödinger, LLC.; <u>PyMOL|</u> pymol.org).

### 2.7. GlyT1 inhibitor assay

ALX5407, a selective human GlyT1 inhibitor(Atkinson *et al*., 2001), was used for an inhibitor assay to functionally disrupt GlyT1 activity (i.e., PFOS-GlyT1 interaction). A dose-response pilot study with ALX5407 in DMSO found 10 nM to be the maximal concentration not affecting HepG2/CA3 cell viability after 6 days consistent with a previous study which reported 3 nM as the EC50 of the inhibitor(Atkinson *et al*., 2001). To assess the effect of GlyT1 inhibition on PFOS cytotoxicity, HepG2/C3A cells were plated into a 96-well plate at a density of 12,000 cells per well a day before exposure. ALX5407 (10 nM) was added to triplicate sets of wells with corresponding DMSO controls and different concentrations of PFOS (0, 75, 150, and 300 µM) at the same time for 6 days. ATP concentration in each well was determined by CellTiter Glo2.0 assay as in 2.1 above. Statistical significance evaluation was done by t-test using GraphPad Prism (version 10.1.0)

### 2.8. Functional enrichment analysis

KEGG gene set enrichment analysis (Log_2_FC values were included) was conducted with sensitive and resistant candidate genes to PFOS via clusterProfiler option using SRPLOT software(Tang *et al*., 2023). STRING database(Szklarczyk *et al*., 2023) was used to cluster protein products of candidate genes that confer sensitivity or resistance to PFOS using default settings (accessed on Mar 13th, 2025). We annotated each cluster of two or more gene products based on Gene Ontology-Biological pathway and KEGG pathway as available within STRING.

### 2.9. Gene-Disease outcome association analysis

Statistically significant candidate genes (189 PFOS-sensitive genes and 151 PFOS-resistant genes) were used to perform over- and under-representation analyses using the CTD(Davis *et al*., 2025) accessed on Jan 30th, 2025. The proportion of PFOS-sensitive and PFOS-resistant genes linked to each Disease was compared to that of equally sized random sets of genes. Disease outcomes with a proportion above 0.001 in our significant candidate gene sets were considered for further analysis. Z-scores were calculated by subtracting our observed value with the mean proportion for each Disease outcomes across 1,000 random sets and divided by the standard deviation of the random sets. *p*-values were obtained using a standard normal distribution, and 0.05 was used as the significance threshold.

### 2.10. G2P-SCAN analysis: Conserved Reactome pathways analysis among other model species

The Genes-to-Pathways Species Conservation Analysis (G2P-SCAN)(Rivetti *et al*., 2023) R package was run using the latest source code for the G2P-SCAN library that is available on GitHub (https://github.com/seacunilever/G2P-SCAN). Based on the gene input, the G2P-SCAN outputs the overall analysis of orthology and functional families to substantiate the identification of conservation and susceptibility at the Reactome pathway level.

## 3. Results

### 3.1 CRISPR screens identified functional genetic components modulating PFOS-induced cytotoxicity

The overall design of this study is illustrated in Fig. 1A. We first identified a dose of PFOS which partially inhibited cellular proliferation of HepG2/C3A over the short-term period of exposure. We determined the IC_25_ for 6 days exposure to be 192 µM for PFOS (Fig. S1), which was further refined to be 170 µM after a pre-screen time-course treatment for the subsequent CRISPR screens. After quality control sequencing of the MinLibCas9 library (Fig. S2), we used the library to generate a mutant pool of knockouts in HepG2/C3A at ∼400X coverage of each sgRNA. After expansion of the mutant pool, the pool was either maintained (Control) or exposed to PFOS (Exposed) at the IC_25_ concentrations for 18 days (approximately 10 doublings). We utilized the IC_25_ of PFOS to identify genes which increase sensitivity to PFOS when disrupted (corresponding sgRNAs depleted in the exposed mutant pool) as well as genes which increase resistance to PFOS when disrupted (corresponding sgRNAs enriched in the exposed mutant pool). We used MAGeCK to identify candidate sgRNAs and genes which when disrupted either increase sensitivity to PFOS or increase resistance to PFOS with the threshold of a *p*-value < 0.01 and Log_2_FC < -0.6 or Log_2_FC > 0.6 (Fig. 1B). At this threshold, we identified a total of 340 candidate genes (189 genes increased sensitivity when disrupted; 151 genes increased resistance when disrupted) to PFOS exposure (a list of the candidate genes in Table S4) (Fig. 1B).

**Fig. 1.**
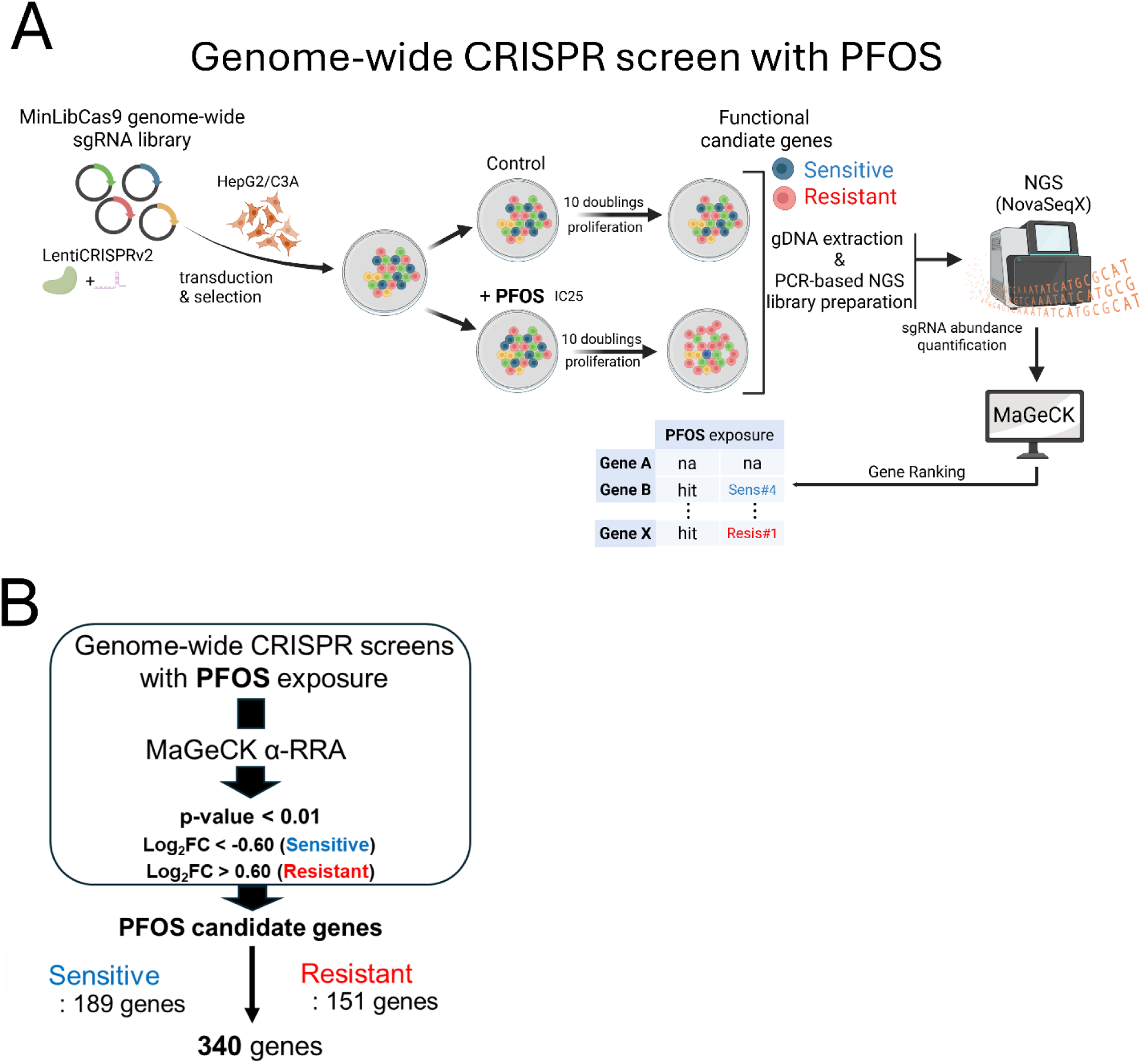
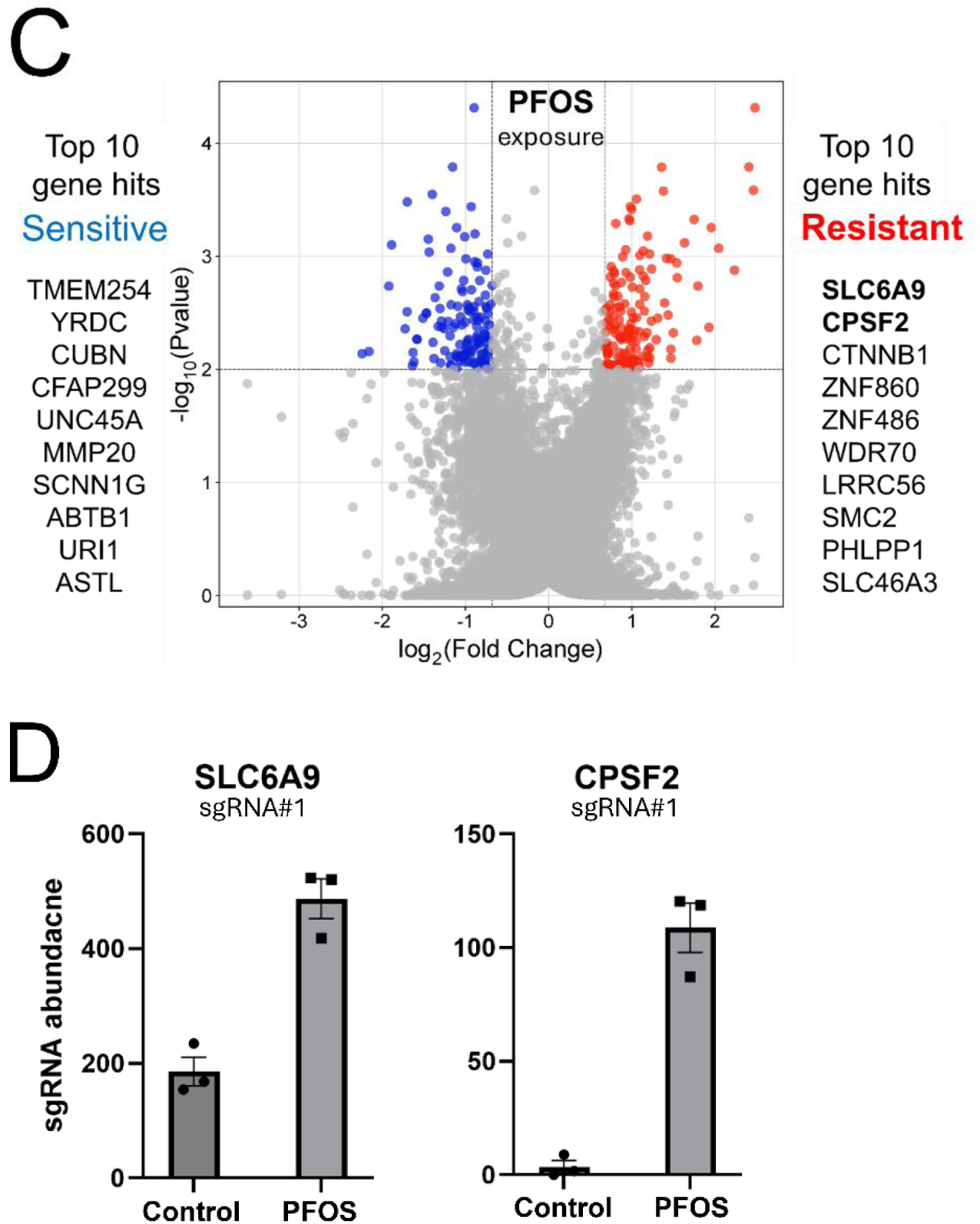
Genome-wide CRIPSR screens to identify functional genetic modulators of PFOS-induced cellular toxicity. The CRISPR screening workflow employed in the study is shown in panel (A), while panel (B) illustrates the bioinformatics analysis and specific criteria used to identify statistically significant candidate genes from the screens. The volcano plot (C) displays candidate genes that modulate PFOS cellular toxicity with different colors. Blue dots represent PFOS-sensitive genes and red dots represent PFOS-resistant genes. The top 10 sensitive and resistant genes are also listed next to the plot. These top candidate genes reflected the most significant ranks from MaGeCK analysis (α-RRA) based on the criteria (B) applied. The x-axis represents Log_2_ FC (fold change), and the y-axis shows -Log10 (*p*-value). Panel (D) shows the representative individual sgRNA abundance results of the top 2 resistant candidate genes (sgRNA#1 for SLC6A9 and CPSF2) from the screens.

The Log_2_FC, *p*-value, and known or predicted functions (abbreviated STRING annotation for corresponding gene product) of the top ranked candidate genes identified in our CRISPR screens are shown in Table 1. The top ten candidate genes that increased resistance to PFOS when disrupted are *SLC6A9, CPSF2, CTNNB1, ZNF860, ZNF486, WDR70, LRRC56, SMC2, PHLPP1,* and *SLC46A3* (Fig. 1C). Similarly, the top ten candidate genes that increased sensitivity to PFOS exposure when disrupted were *TMEM254, YRDC, CUBN, CFAP299, UNC45A, MMP20, SCNN1G, ABTB1, URI1,* and *ASTL*. Analysis using a different algorithm, PinAPL-Py(Rockett, 2002), identified a similar set of candidates particularly among the significantly ranked genes (all top sensitive and resistant genes were identified as significant except for *ASTL* gene) (Fig. S3). We selected the two most significant candidate genes identified in both analysis methods as top-ranked genes that increase resistance to PFOS exposure, *SLC6A9* and *CPSF2* for further investigation. Individual sgRNA abundance of the selected genes in our screening data (normalized sgRNA counts for each gene in our CRISPR screens) showed an increase in abundance upon PFOS exposure (Fig. 1D).

**Table 1.**
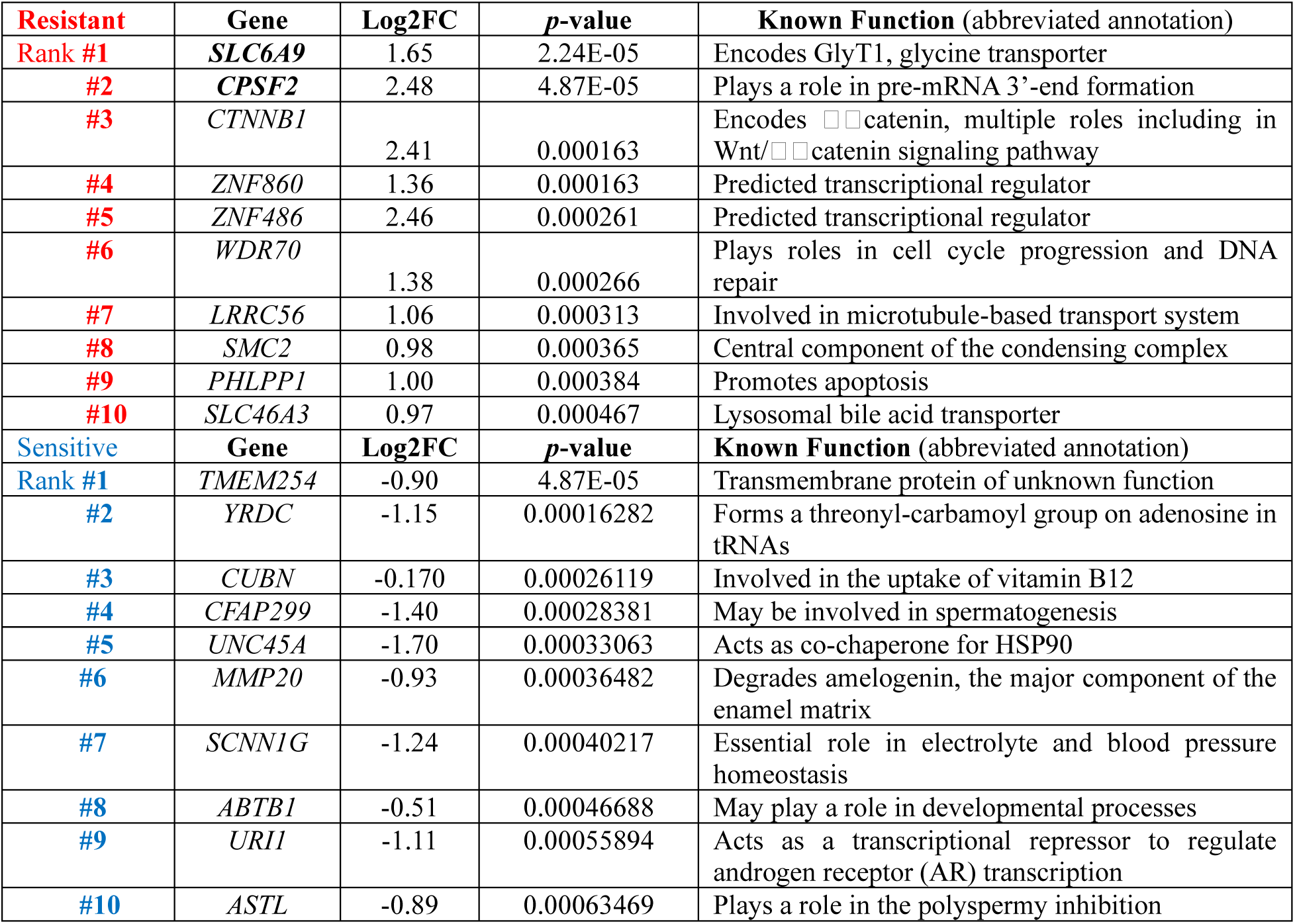
Summary of top 10 genes ranked by MaGeCK αRRA which increase resistance to PFOS exposure when disrupted (Red category) and top 10 genes ranked by MaGeCK αRRA which increased sensitivity to PFOS when disrupted (Blue category). The Log_2_FC and *p*-value calculated from MaGeCK analysis is shown for each gene, with brief functional annotation of the corresponding gene product from STRING database(Szklarczyk *et al*., 2023) (<u>STRING: functional protein association networks</u>).

### 3.2 Functional validation of selected candidate genes supports a role for *SLC6A9* and *CPSF2* during PFOS exposure

To validate the selected candidate genes identified as being resistant to PFOS in our screens, we performed single gene CRISPR/Cas9 targeting of *SLC6A9* and *CPSF2*. Consistent with the results of the CRISPR screens, the cell pools with single gene knockouts (KOs) validated at the gene and mRNA levels (Fig. S4 and Fig. 2A) exhibited dose-response cellular resistance to PFOS exposure for 6 days with improved cell viability compared to wild-type HepG2/C3A cells (Fig. 2B and Fig. 2C). Disruption of the *SLC6A9* in HepG2/C3A cells resulted in significantly increased resistance to PFOS exposure at 150 µM (*p*= 0.00013) and 300 µM (*p*= 0.00096) (Fig. 2B) and disruption of *CPSF2* similarly showed significantly enhanced cell viability upon PFOS exposure at 150 µM (*p*= 0.0002) and 300 µM (*p*= 0.00149) (Fig. 2C). The results support a functional role for the corresponding gene products in modulating PFOS-induced cytotoxicity.

**Fig. 2.**
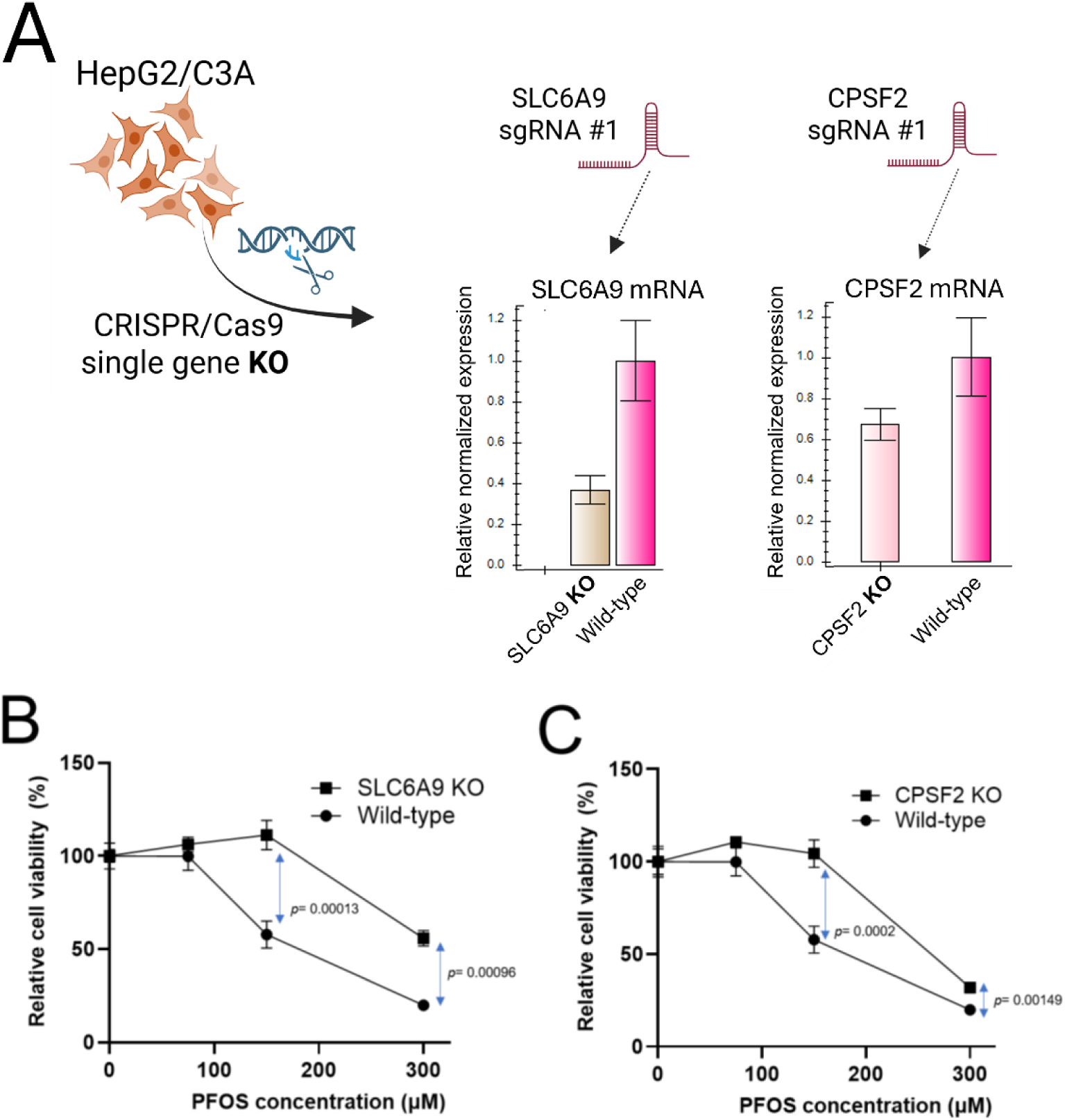
Targeted validation of candidate genes identified from the PFOS CRISPR screens. For the top 2 candidate genes resistant to PFOS exposure, *SLC6A9* and *CPSF2*, individual knockouts (KOs) of each gene are generated in HepG2/C3A cells by single gene CRISPR/Cas9 with the effect of each sgRNA#1, targeting each gene, on corresponding mRNA levels is shown (A). Using these KO cells, cell viability assays were performed and compared to wild-type HepG2/C3A cells for validation. Cell viability for 6-day exposure to PFOS was used to measure the relative chemical susceptibility (B and C). Three replicates were used for cell viability assays.

### 3.3 Inhibitor assay and molecular docking study suggest PFOS interacts with GlyT1 encoded by *SLC6A9*

Our screening and validation results showed that the functional disruption of *SLC6A9*, which encodes Glycine Transporter 1 (GlyT1), increased cellular resistance to PFOS exposure. Disruption of a cellular transporter resulting in increased resistance to a xenobiotic is consistent with a potential role in transport of the xenobiotic(Klaassen, 2002; George *et al*., 2017). We next tested whether ALX5407, a selective irreversible inhibitor of human GlyT1(Atkinson *et al*., 2001), induces cellular resistance to PFOS exposure (Fig. 3A). Indeed, HepG2/C3A cells treated with ALX5407 showed significantly increased resistance (*p*= 0.025) to PFOS exposure at 300 µM for 6 days in the presence of 10 nM ALX5407 (Fig. 3A), consistent with a functional role for GlyT1 in modulating PFOS toxicity. No interaction with or role of GlyT1 in PFOS transport has been previously reported to our knowledge, although other solute carrier transporters (SLCs), including sodium/taurocholate co-transporting polypeptide (NTCP) and organic anion transporter 4 (OAT4), have been reported to transport PFOS in several cell types(Vujic, Ferguson and Brouwer, 2024). We noted that Bitopertin, a known inhibitor of GlyT1, showed possible structural similarity to PFOS with a sulfonyl group (SO_2_CH_3_) and multiple trifluoromethyl groups (CF_3_) (Fig. 3B). Previous studies had carried out molecular docking studies between Bitopertin and GlyT1 and GylT2 (encoded by *SLC6A5*)(de Carvalho *et al*., 2024), and thus we employed molecular docking to predict potential intermolecular contacts between PFOS and the GlyT1 transporter as well as the closely related GlyT2. We hypothesized that PFOS might interact with amino acid side chains in the central regions of the GlyT1 and GlyT2 transporter similar to as seen with Bitopertin(de Carvalho *et al*., 2024). Indeed, docking simulations using AutoDock Vina predicted PFOS binding to a conserved site within the GlyT1 and GlyT2 transporters. In GlyT1, PFOS was predicted to form hydrogen bonds with residues Y196, Y370, S371, and T472, (ΔG value -9.5 kcal/mol) and in GlyT2, predicted hydrogen bonding occurs with residues at similar corresponding positions, Y289, F478, S481, and T580 (Fig. 3C). ΔG scores were -9.5 kcal/mol and -9.4 kcal/mol for PFOS interaction with GlyT1 and GlyT2, respectively, as compared to predicted ΔG of -3.89 for glycine interaction with GlyT1 at this site and ΔG of -4.3 for glycine interaction with GlyT2 at this site (Fig. 3C). Based on the potential interaction of PFOS with GlyT1 predicted by molecular docking simulation (potential substrate or inhibitor), inhibition assay (enzymatic inhibition), and KO validation (genetic disruption), we suggest that GlyT1 (*SLC6A9*) modulates PFOS cellular toxicity through a direct interaction with PFOS, but we cannot differentiate between a role for PFOS as an GlyT1 inhibitor or as a substrate. (B) Structural similarity between another GlyT1 inhibitor, Bitopertin, and PFOS. Red colors indicate putative structural similarity to PFOS with a sulfonyl group (SO_2_CH_3_) and multiple trifluoromethyl groups (CF_3_). (C) Molecular docking results present predicted intermolecular contact between GlyT1 (encoded by *SLC6A9*), GlyT2 (encoded by *SLC6A5*) and PFOS. PFOS is shown as sticks, with yellow for carbon, orange for sulfur, red for oxygen, and light blue for hydrogen. In the upper part, GlyT1 was depicted as a copper-colored ribbon diagram. In the lower part, GlyT2 was depicted as a green ribbon diagram. Residues involved in hydrogen bonding with PFOS were shown as stick models, with violet for carbon, red for oxygen, and blue for nitrogen. Residues that were predicted to form hydrogen bonds with PFOS are shown as stick models, with cyan representing carbon, red representing oxygen, and blue representing nitrogen.

**Fig. 3.**
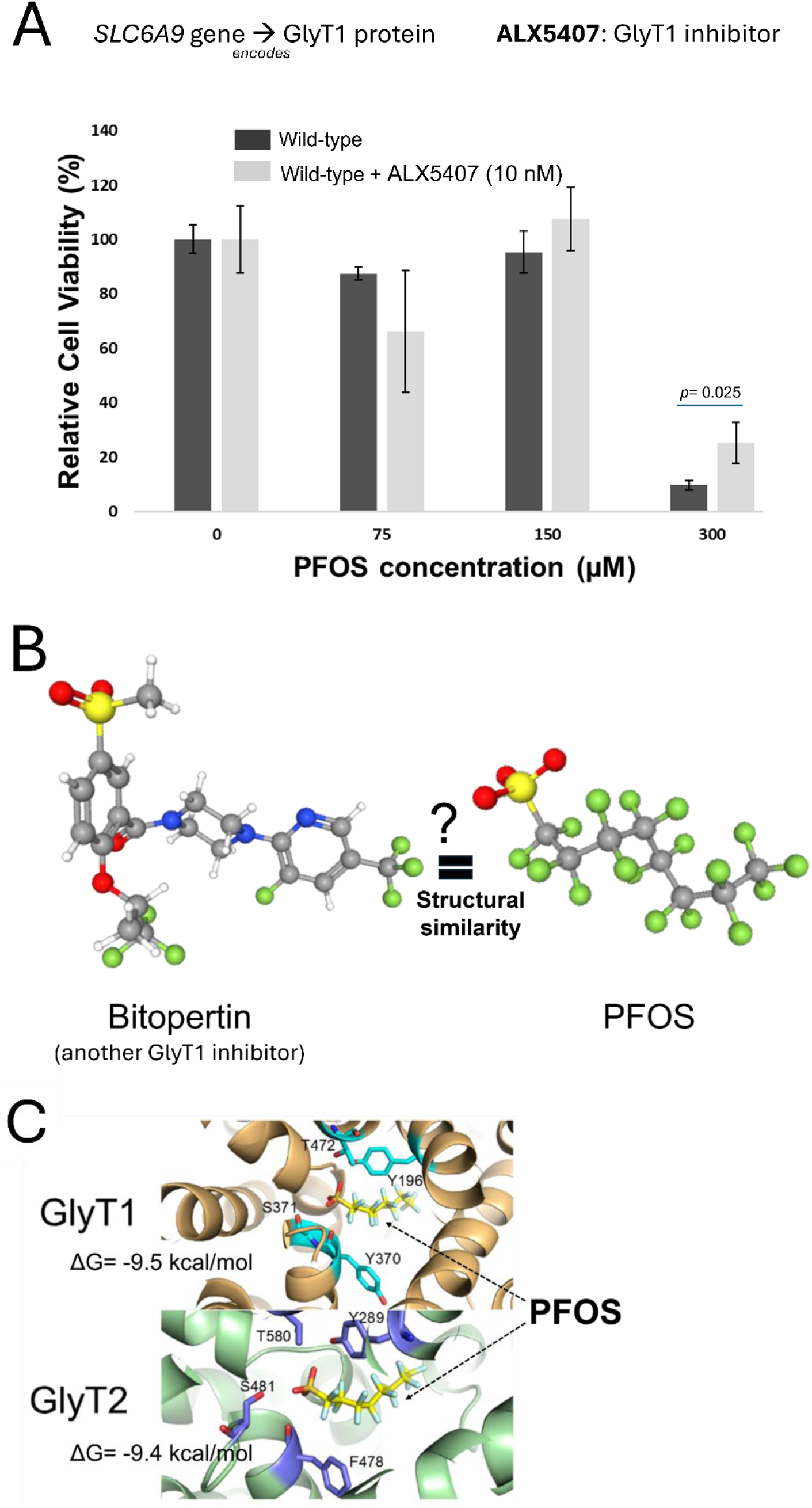
Further analysis of the top resistant candidate gene for PFOS toxicity, *SLC6A9,* and predicted binding between GlyT1 (encoded by *SLC6A9*) and PFOS. (A) Enzymatic activity inhibition of GlyT1 by treatment of ALX5407 and its functional effect on PFOS-induced cellular toxicity. Cell viability of HepG2/C3A cells exposed to PFOS (0, 75, 150, 300 µM) when treated with ALX5407 (10 nM), a selective GlyT1 inhibitor for 6 days. Three replicates were used for cell viability assays. Statistical significance evaluation was done by t-test using GraphPad Prism (version 10.1.0).

### 3.3 Gene-Disease outcome association analysis using the Comparative Toxicogenomic Database indicated cancer (neoplasm, carcinoma) and liver disease as potential adverse outcomes of PFOS toxicity

To survey the potential association with adverse outcomes (disease phenotypes) of our significant candidate genes, we performed a custom Gene-Disease outcome association analysis using the CTD (Fig. 4A). We identified disease terms over-represented in the PFOS-sensitive and PFOS-resistant genes. The top 15 over-represented diseases for the PFOS-sensitive gene set as compared to randomly sampled gene sets of the same size are shown in Fig. 4B. 8 of the 15 top over-represented diseases in PFOS-sensitive genes included terms related to cancer such as Pancreatic Neoplasms, Adenocarcinoma, Prostatic Neoplasms, and Liver Neoplasms (Fig. 4B). In contrast, the analysis of the PFOS-resistant genes found 6 out of the top 15 associated phenotypes are related to liver disease, including liver failure, cholestasis, and liver cirrhosis (Fig. 5A).

**Fig. 4.**
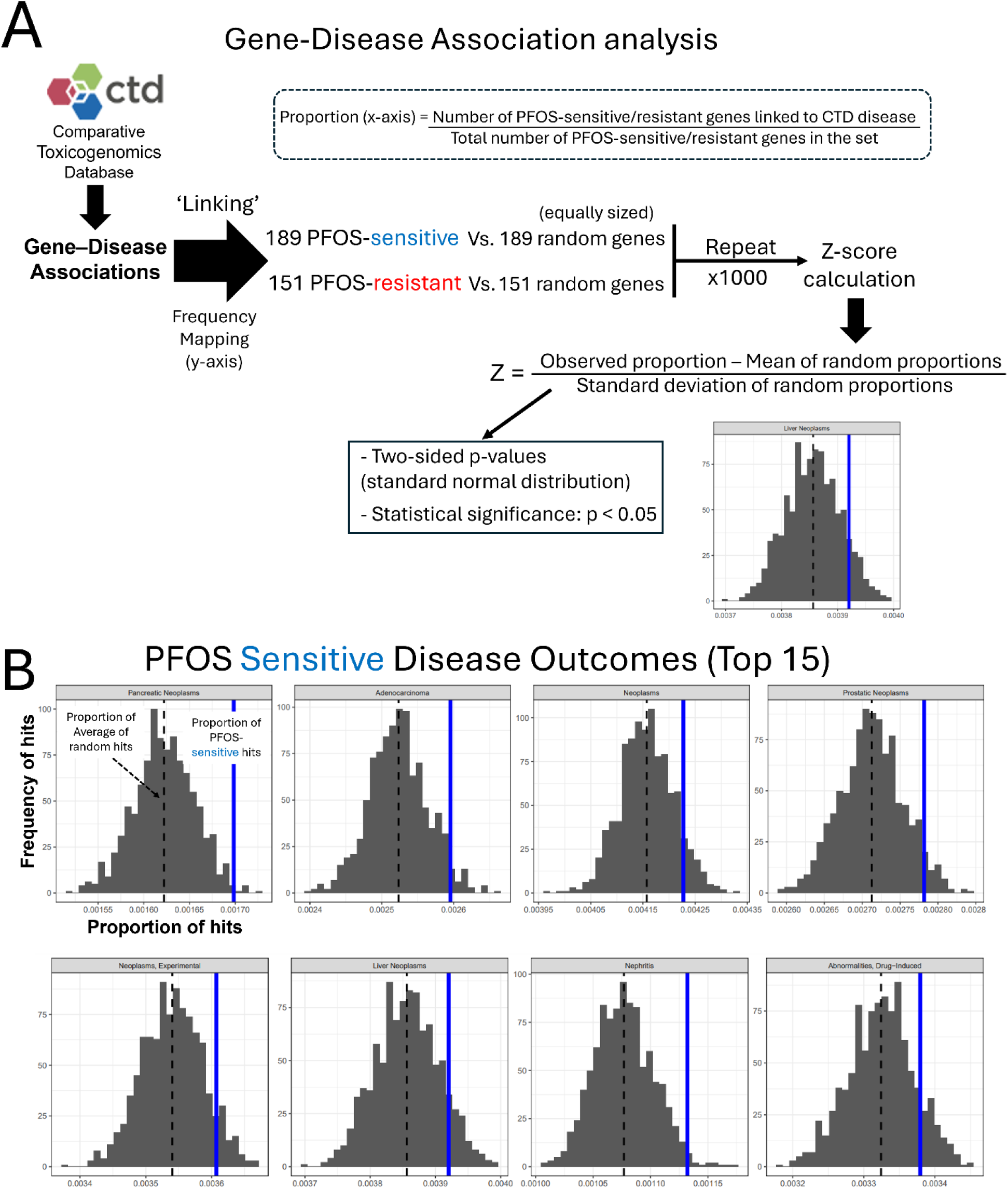
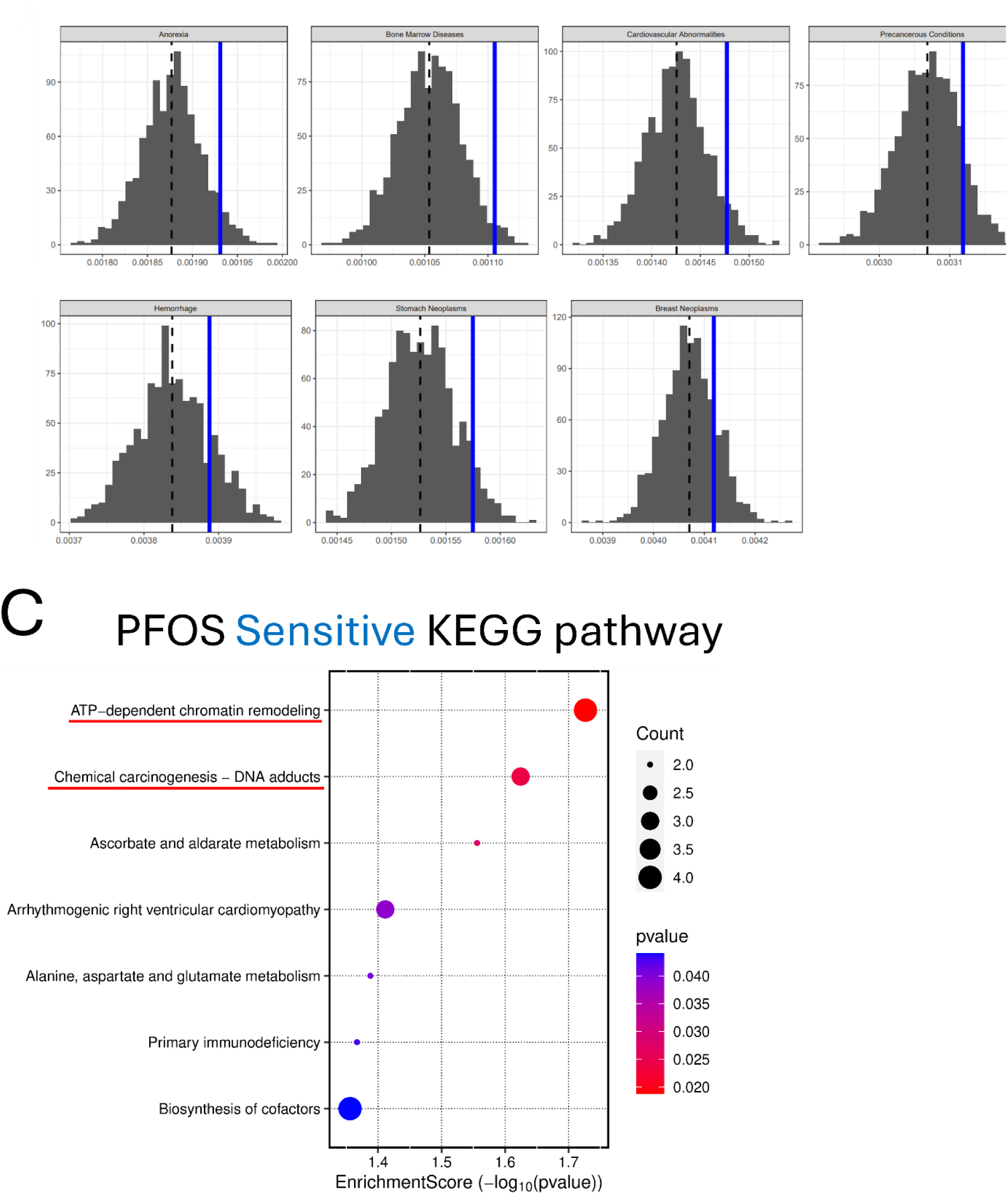
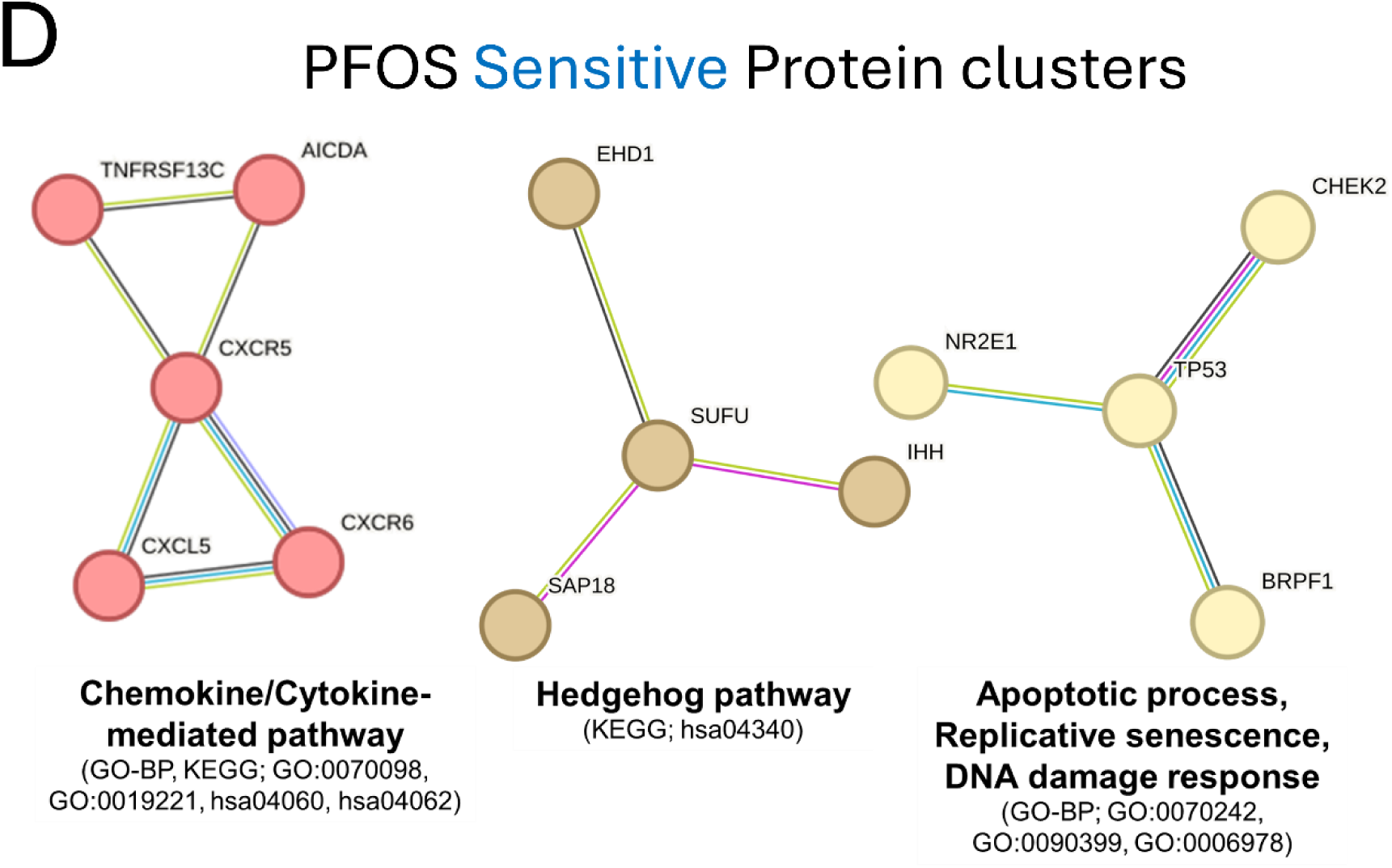
Functional enrichment analyses of PFOS-sensitive genes. (A) Schematic overview of the Gene-Disease outcome analysis workflow. (B) Results of the top 15 significant disease outcomes over-represented in PFOS-sensitive genes are presented. In the histograms, the x-axis indicates the proportion of Disease hits with the dashed black line representing the mean of the random sets shown in gray, and our observed proportion represented by the solid blue line. The y-axis indicates the frequency of the proportion in the randomly sampled sets. The disease categories are displayed at the top of each plot. (C) KEGG pathway enrichment result of PFOS-sensitive genes. (D) STRING clusters of the functional protein interactions enriched in PFOS-sensitive genes.

**Fig. 5.**
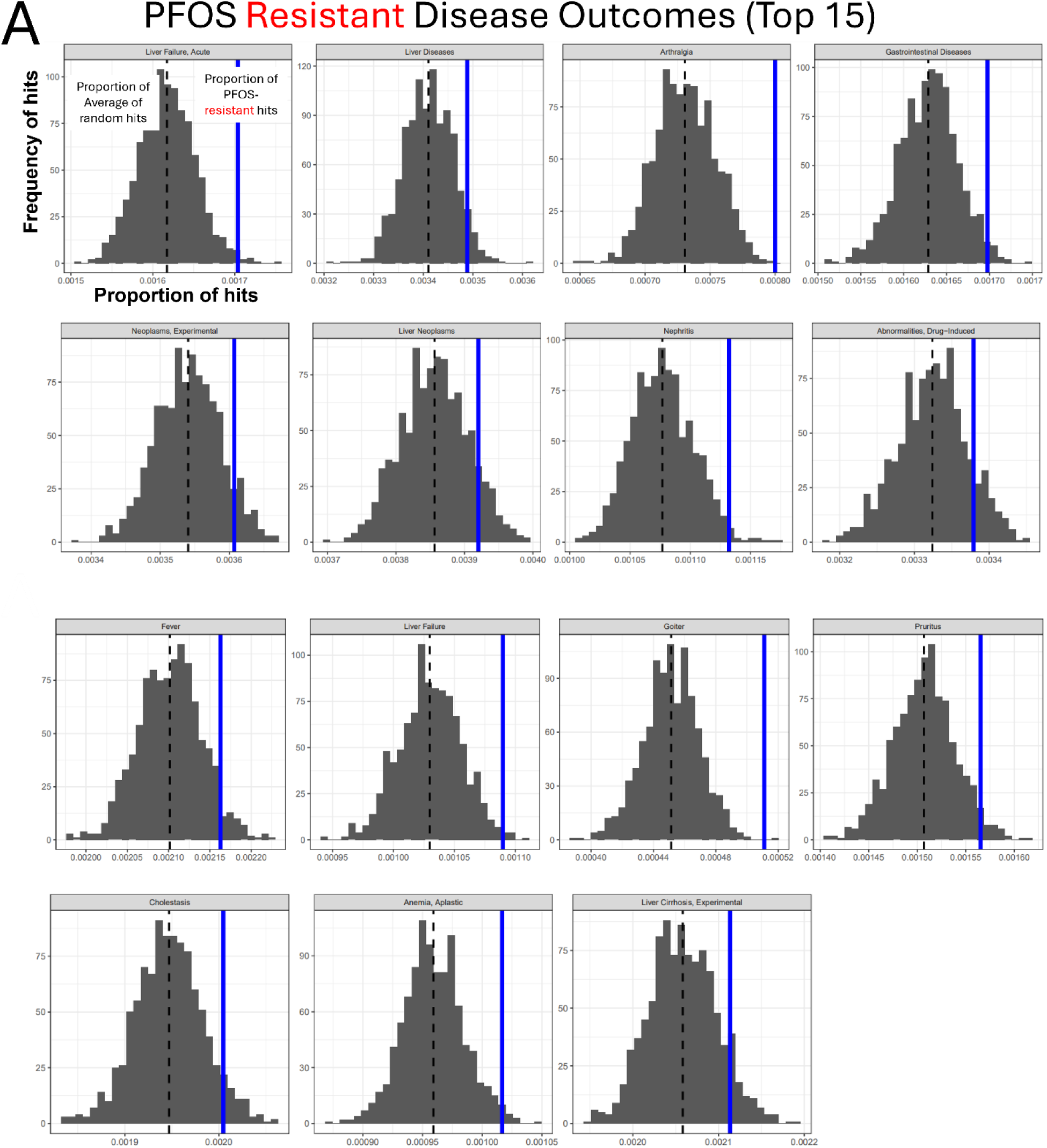
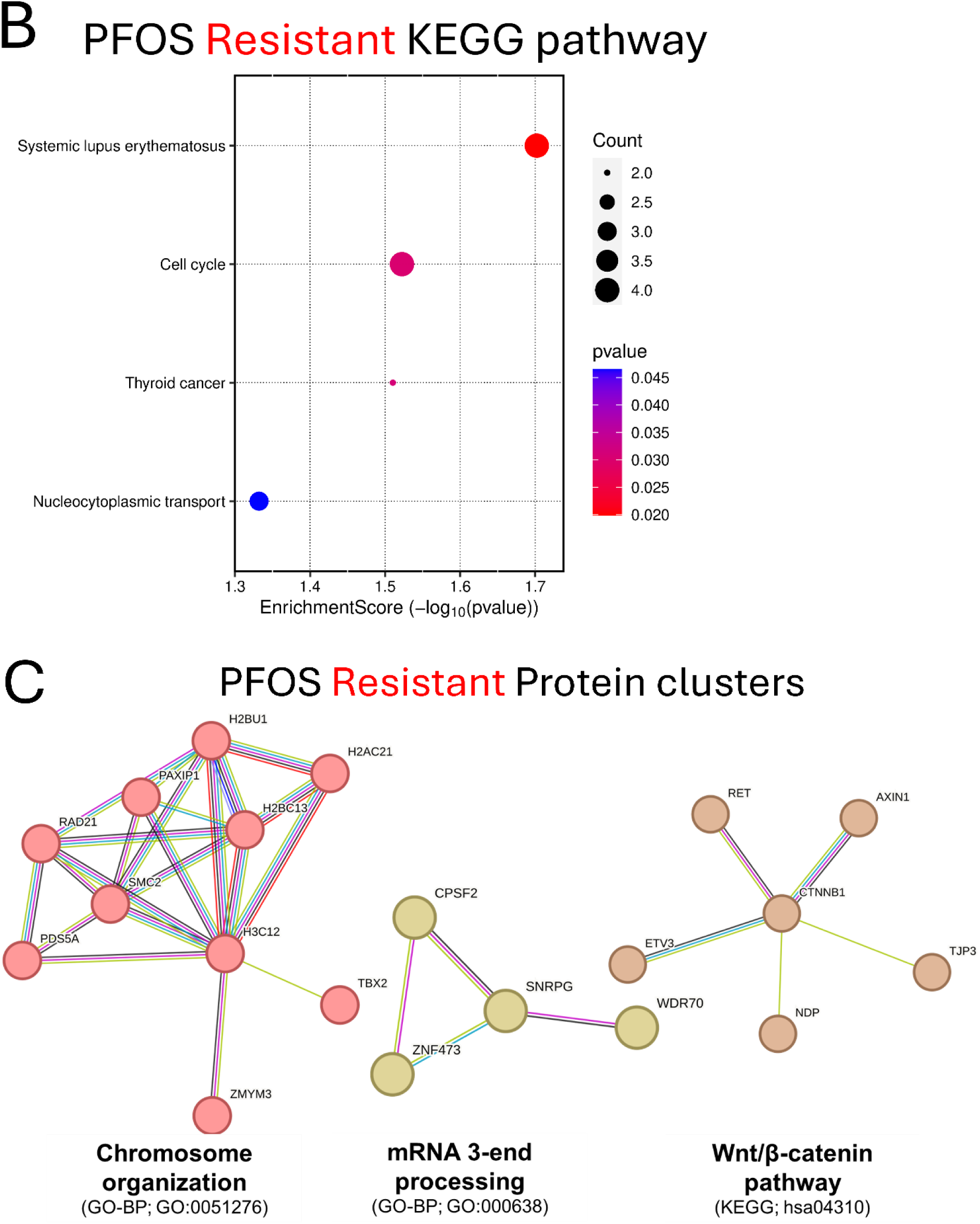
Functional enrichment analyses of PFOS-resistant genes. (A) Results of the top 15 significant disease outcomes over-represented in PFOS-resistant genes. In the histograms, the x-axis indicates the proportion of Disease hits with the dashed black line representing the mean of the random sets shown in gray, and our observed proportion represented by the solid blue line. The y-axis indicates the frequency of the proportion in the randomly sampled sets. The disease categories are displayed at the top of each plot. (B) KEGG pathway enrichment result of PFOS-resistant genes. (C) STRING clusters of the functional protein interactions enriched in PFOS-resistant genes.

### 3.4 Enrichment analyses identified potential cellular pathways modulating PFOS toxicity

Next, we carried out KEGG and STRING analysis on candidate genes (sensitive and resistant genes) identified in the screens to gain insight into potential involved pathways and to explore functional interactions modulating PFOS-induced cytotoxicity (Fig. 4C, 4D, 5B, and 5C). KEGG gene set enrichment analysis revealed that ATP-dependent chromatin remodeling and chemical carcinogenesis-DNA adducts related pathways are among the most significantly enriched pathways in PFOS-sensitive genes (Fig. 4C), consistent with the results of CTD Gene-Disease outcome association analysis which found cancer-related phenotypes being significantly over-represented (Fig. 4B). Functionally annotated STRING clusters of the PFOS-sensitive genes included Chemokine and Cytokine-mediated signaling pathway (*CXCR5, CXCL5, CXCR6, AICDA, TNFRSF13C*), the Hedgehog signaling pathway (*SUFU, IHH, SAP18, EHD1*), and cell cycle-related pathway such as Apoptotic process, Replicative senescence, and DNA damage response (*TP53, NR2E1, CHEK2, BRPF1*) (Fig. 4D) as well as Purine biosynthesis, Nuclear pore complex disassembly, Synaptic membrane adhesion, and others (Fig. S5A). KEGG analysis of PFOS-resistant genes identified Systemic lupus erythematosus, Cell cycle, and others as significantly enriched pathways (Fig. 5B). Functionally annotated clusters identified by STRING analysis of the PFOS-resistant genes included Chromosome organization (*H3C12, TBX2, ZMYM3, H2BC13, H2AC21, SMC2, PDS5A, RAD21, PAXIP1, H2BU1*), mRNA 3-end processing (*SNRPG, CPSF2, WDR70, ZNF473*), the Wnt signaling pathway (*CTNNB1, AXIN1, TJP3, RET, ETV3, NDP*) (Fig. 5C) as well as several others including Ionotropic glutamate receptor complex and Regulation of oxidative phosphorylation (Fig. S5B).

### 3.5 G2P-SCAN analysis provides cross-species applicability of adverse effects using gene targets of PFOS-induced toxicity

To provide insight into potential cross-species PFOS toxicity based on our human cell-based functional data, we conducted an *in silico* cross-species pathway prediction using G2P-SCAN. The G2P-SCAN analyzes data from various databases, integrating gene orthologs, protein families, entities, and reactions linked to human genes and respective pathways across six relevant model species including rat (*Mus musculus*), mouse (*Rattus norvegicus*), zebrafish (*Danio rerio*), fruit fly (*Drosophila melanogaster*), nematode (*Caenorhabditis elegans*), and budding yeast (*Saccharomyces cerevisiae*). We focused on top significant candidate genes including *SLC6A9* and *CPSF2*, the two experimentally validated genes. *SLC6A9* gene was mapped to only a single pathway (i.e., Na+/Cl- dependent neurotransmitter transporters, Fig. 6A) compared to result of *CPSF2* (Fig. 6B) which showed mapping to multiple pathways, including RNA polymerase II transcription termination, Processing of intronless pre-mRNAs, and mRNA 3’-end processing. While *SLC6A9* showed 100 % pathway conservation in mammals, zebrafish and non-vertebrate species showed much lower conservation (zebrafish: 74 %, nematode: 21 %, fruit fly: 32 %, Budding yeast: 0 %) of this pathway. *CPSF2* was highly conserved across vertebrates in the related pathways (83 % to 100 %) whereas we observed lower conservation in other species (39 % to 78 %) with notable exception of fruit fly (93%) in the pathway of ‘transport of mature mRNA derived from an intronless transcript’. Together, these results showcased that functionally identified gene targets responsible for a certain chemical-induced toxicity can be used to predict a potential role in other species under the same environmental exposure and that conservation levels of gene targets and related pathways may be different across the species.

**Fig. 6.**
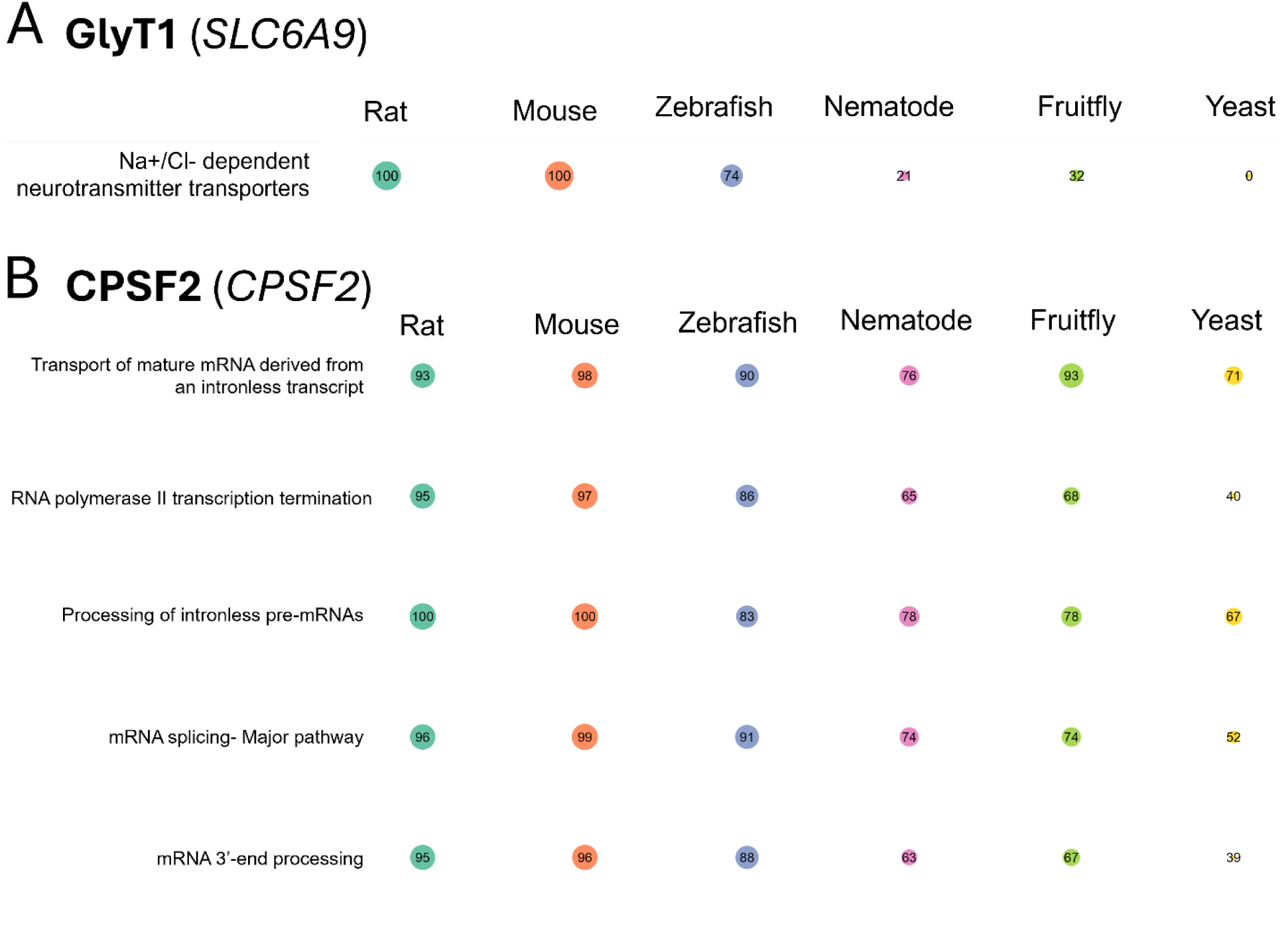
The predicted conservation percentage for each species compared to the human ortholog based on the Genes-to-Pathways Species Conservation Analysis (G2P-SCAN) output, using GlyT1 (*SLC6A9*) (A) and CPSF2 (*CPSF2*) (B) as inputs, is shown. The individual conservation level related to each human gene input was analyzed using G2P-SCAN to determine the level of conservation across six model species presented by Reactome pathways. The size of circle is proportional to the percentage of conservation. The color of circle indicates different species, including Rat, Mouse, Zebrafish, Nematode, Fruit fly, and Yeast.

## 4. Discussion

Extensive epidemiological studies as well as controlled exposure studies in multiple species provide support for PFOS exposure being associated with adverse outcomes on liver function(Fenton *et al*., 2021; Jane L Espartero *et al*., 2022) while the mechanisms underlying adverse outcomes remain controversial. Several potential mechanisms may play a role such as Peroxisome proliferator-activated receptor alpha (PPARα) activation(Azhagiya Singam *et al*., 2024), but definitive causal connections remain elusive. To provide insights into the functional effects of PFOS on the human liver, we carried out a genome-wide CRISPR screen with PFOS in this study. We identified multiple candidate genes that when disrupted modulate the cellular toxicity of PFOS and among these candidates, two selected genes (*SLC6A9* and *CPSF2*) were experimentally validated, recapitulating the screening results. We further identified potential associated phenotypic outcomes (i.e., diseases), cellular pathways, and molecular networks potentially involved in PFOS toxicity using the PFOS candidate genes, which together suggest previously unrecognized potential mechanisms of PFOS-induced adverse effects.

Identification of the GlyT1 as a candidate gene functionally modulating PFOS-induced cellular toxicity, which transports glycine into multiple cells and tissues (including the hepatocyte in the liver(Howard *et al*., 2010; Wei *et al*., 2024)), suggests a previously unknown connection between this amino acid transporter and PFOS. Several previous studies have noted increased mRNA expression level of *SLC6A9* associated with PFOS exposure in mice(Pfohl *et al*., 2021; Deng *et al*., 2022) and *in vitro*(Louisse *et al*., 2023) but the effect of genetic disruption of the gene on PFOS toxicity hasn’t been studied. We considered the hypothesis that PFOS may directly interact with GlyT1 based on a previous study which carried out molecular docking of a GlyT1 inhibitor, Bitopertin, with structural features reminiscent of PFOS (Fig. 3B). Our molecular docking studies of PFOS with structural models of GlyT1 and the closely related GlyT2 unexpectedly predicted considerable affinity of PFOS for both GlyT1 and GlyT2 in the same region with the known inhibitor, with a predicted G values exceeding the interaction with glycine alone (Fig. 3C). This potential interaction was further supported by chemical disruption of GlyT1 enzymatic activity with another selective GlyT1 inhibitor, ALX5407, which also conferred resistance to PFOS exposure (Fig. 3A). Together, molecular docking and enzymatic inhibition studies with SLC6A9 KO validation suggest that functional disruption of GlyT1 confers increased cellular resistance to PFOS exposure possibly via disrupted or reduced interaction between GlyT1 and PFOS. However, further follow-up studies such as PFOS and glycine (natural substrate) competitive uptake studies would be needed to confirm direct interaction. A recent review for the role of xenobiotic transporters expressed in the liver in PFAS transport noted that most of functional studies have relied on heterologous expression of transporters in cell lines(Vujic, Ferguson and Brouwer, 2024). Indeed, the Na^+^-taurocholate co-transporting polypeptide (NTCP) and several members of the organic anion transporting polypeptides (OATP) have been identified to play roles in PFOS uptake and reabsorption(Louisse *et al*., 2024). However, there remains uncertainty about the physiological role of the NTCP and OATP transporters, particularly in the liver. We did not find any of these previously identified transporters in our genetic screens, which indicates that they may not play primary roles in HepG2/C3A, or alternatively, functional redundancy may exist (e.g., multiple transporters could be involved and loss of single one may not alter the cellular response). In sum, our results suggest a possible role of GlyT1 in modulating PFOS cytotoxicity which has not been previously documented to our knowledge. While our studies primarily suggest a direct interaction of PFOS with GlyT1 underlies its role in PFOS-induced toxicity, further study will be needed to determine if it represents a substrate of GlyT1. Disruption of *CPSF2* resulted in increased resistance to PFOS toxicity as well (Fig. 2A and 2C). *CPSF2* encodes the CPSF2 protein which mediates 3’ end pre-mRNA processing, including poly-adenylation and histone mRNA 3’ end processing(Dominski and Tong, 2021). The exact role of *CPSF2* remains controversial(Liu and Manley, 2024) and evidence for involvement of CPSF2 in mediating PFOS toxicity is limited. While several studies have identified histone epigenetic modifications associated with PFOS exposure(Kim, Thapar and Brooks, 2021; Li *et al*., 2024), direct effects on histone mRNA processing have not been noted to our knowledge. Interestingly, a genome-wide association study (GWAS) of perfluorooctanoic acid (PFOA) *in vitro* toxicity identified the *CPSF2* locus as a candidate modulator of cytotoxicity(O’Shea *et al*., 2011). Transcriptional studies in zebrafish done by Dasgupta et al. and Rericha et al. with perfluorooctane sulfonamide (PFOSA), a precursor of PFOS, reported conflicting results with increased^70^ or decreased^71^ *CPSF2* mRNA expression, respectively. *CPSF2* was identified as hub gene involved in lipid metabolism from weighted gene co-expression network analysis on RNA-seq data of bovine samples(Wang *et al*., 2022) potentially in line with lipid metabolism disruption which is suggested as one of molecular mechanisms of PFAS toxicity(Evans *et al*., 2022).

We used the CTD(Davis *et al*., 2025) to identify potential correlations between the candidate genes identified in our screens and associated disease phenotypes curated in the CTD. Surprisingly, the PFOS-sensitive genes showed over-representation in cancer-related terms, including Pancreatic neoplasms, Adenocarcinoma, Prostatic neoplasms, and Liver neoplasms among the top over-represented diseases. The adverse outcomes of PFAS exposure in general as well as PFOS specifically in the risk of developing different forms of cancer, including of the liver, kidney, thyroid, and blood remains uncertain(Steenland and Winquist, 2021; Gerwen *et al*., 2023; Rhee *et al*., 2023; Winquist *et al*., 2023). The International Agency for Research on Cancer (IARC) classified PFOS as a Group 2B carcinogen (possibly carcinogenic) in Dec 2023(Zahm *et al*., 2024;

*IARC Monographs evaluate the carcinogenicity of perfluorooctanoic acid (PFOA) and perfluorooctanesulfonic acid (PFOS)*, no date). Several studies of PFOS exposure in rodents find evidence for hepatotoxicity, liver dysfunction(Bjork, Butenhoff and Wallace, 2011; Costello *et al*., 2022), and an early study in a rat model chronically exposed to PFOS observed significant increases in hepatocellular adenoma(Butenhoff *et al*., 2012). Furthermore, KEGG and STRING enrichment analyses identified several cancer-related pathways and protein networks, respectively. The ATP-dependent chromatin remodeling (Fig. 4C) pathway can play a role as a key driver of cancer in some contexts, including by promoting the abnormal gene expression that maintains cancer status(Mayes *et al*., 2014) and/or by altering the accessibility of DNA in the chromatin to DNA-binding factors, which regulates DNA replication and DNA repair(Zhang *et al*., 2011). The Chemical carcinogenesis-DNA adducts (Fig. 4C) pathway is also closely related to cancer as the accumulation of DNA damage or DNA adduct formation is a necessary process for tumor induction and progression and increases cancer risk(Poirier, 2012). Consistent with these results, STRING network analysis of the gene products responsible for PFOS sensitivity identified a cluster of genes related to cell cycle processes, including apoptosis, DNA replication, and the DNA damage response. In marine vertebrates and bacteria(Liu *et al*., 2016; Otero-Sabio *et al*., 2022), cell cycle alterations and DNA damage response has been postulated as potential mechanisms of PFOS-induced toxicity. The enrichment of a DNA damage response protein network in the PFOS-sensitive genes is consistent with the cancer-related phenotype association from the CTD analysis.

Another protein interaction cluster comprised of CXCR5, CXCL5, CXCR6, AICDA, TNFRSF13C involved Chemokine and Cytokine-mediated signaling pathways. Chemokines are known to play a crucial role in maintaining the innate immune system(Mackay, 2001), and epidemiological studies have linked immune dysfunction with PFOS exposure(Lee *et al*., 2018; Ehrlich *et al*., 2023). A 3D *in vitro* study to examine PFOS effects on cardiac development reported decreased expression of Hedgehog pathway genes, including *HHIP*, *HHATL*, and *LQCE*, as well as an increased expression of *ENPP1*, a Hedgehog repressor, suggesting PFOS exposure impairs the Hedgehog signaling pathway(Davidsen *et al*., 2022).

In contrast, we identified over-representation of liver disease-related terms in our PFOS-resistant genes. In people, several epidemiological studies linked PFOS exposure to elevated levels of alanine aminotransferase (ALT) in serum, a widely used marker of liver damage(Addicks *et al*., 2023; Zhang *et al*., 2023). A rodent study demonstrated that PFOS exposure exacerbates alcohol-induced liver injury, suggesting its potential as a significant risk factor in this phenotype(Ekuban *et al*., 2025). KEGG analysis identified Systemic lupus erythematosus (SLE) and cell cycle as significant pathways enriched for PFOS-resistant genes (Fig. 5B). Interestingly, liver abnormalities defined as higher levels of liver damage markers have been found among newly diagnosed SLE patients(Bessone, Poles and Roma, 2014). Cell cycle was also identified as enriched in KEGG (Fig. 5B) analysis and PFOS exposure *in vitro* was reported to promote hepatic cell proliferation, as evidenced by biomarkers such as nuclear protein Ki67 (Ki67) and topoisomerase 2 alpha (Top2α)(Cui *et al*., 2015). The ‘liver neoplasms’ CTD over-representation (Fig. 5A) could be attributed to cell cycle-related pathway enrichment and corresponding genes (cell cycle: PDS5A, ORC6, RAD21, ZBTB17; Thyroid cancer: CTNNB1, RET; see Table S5) identified from our CRIPSR screens. STRING clusters of PFOS-resistance gene products included protein networks involved in chromosome organization, mRNA 3’ end processing, and the Wnt signaling pathway (Fig. 5C). Among these, the Wnt/b-catenin signaling pathway was linked to PFOS-induced liver damage in *in vitro* study where PFOS causes oxidative stress attributed to activation of the Wnt/b-catenin signaling pathway(Jiajing *et al*., 2024). Moreover, exposure to low concentration of PFOS (10 µM) was reported to alter Wnt signaling in mouse bone cells, suggesting the importance of Wnt/ -catenin signaling pathway in PFOS toxicity mechanisms(Xue *et al*., 2025). Overall, the gene-disease outcome association analysis highlights potential connection between the functional genetic components modulating PFOS toxicity and cancer and liver disease phenotypes.

The One Health approach(Buttke, 2011; Sauvé, 2024) can take advantage of available data in diverse species to extrapolate the effects of a chemical on molecular targets, affected biological pathways, and signaling cascades in one species to molecular targets conserved in other exposure species. We used G2P-SCAN analysis to assess the relevance of our studies in human cells to other species and identify candidate molecular targets conserved in those species(Schumann *et al*., 2024). The cross-species comparison of human gene targets specifically linked to PFOS-mediated chemical toxicity, such as *SLC6A9* and *CPSF2*, enabled the estimation of the relevance of these targets mediated toxicity in other species. While the applicability of such target-based inference of adverse effects depends on the conservation of molecular targets and pathways across species, this cross-species prediction tool can help identify priority species that may be at higher risk through similar mechanisms to an environmental toxicant to which humans are being exposed(Rivetti *et al*., 2023; Schumann *et al*., 2024).

## CRediT author contribution

<u>Chanhee Kim</u>: Conceptualization, Investigation, Methodology, Formal analysis, Visualization, Writing- original draft preparation, <u>Abderrahmane Tagmount</u>: Investigation, Methodology, Formal analysis, Writing- review and editing, <u>Zhaohan Zhu</u>: Methodology, Data analysis, Writing- review and editing, <u>Frances Wilson</u>: Investigation, Methodology, <u>Danmeng Li</u>: Methodology, Formal analysis, Writing- method, review, and editing, <u>David A. Ostrov</u>: Writing- review and editing, <u>W. Brad Barbazuk</u>: Writing- review and editing, <u>Rhonda L. Bacher</u>: Investigation, Methodology, Data analysis, Writing- method, review, and editing. <u>Chris D. Vulpe</u>: Conceptualization, Supervision, Writing- review and editing, funding acquisition.

## Acknowledgements

This study was supported by NIEHS (the grant number is R01ES033625).

## Data availability

The raw and processed CRISPR screen data with the corresponding metadata in this study were deposited into Gene Expression Omnibus (GEO) database repository and available with an accession number of GSE291529.

