## Supplementary Figures for "Identification of Functional Genetic Components Modulating Toxicity Response to PFOS using Genome-wide CRISPR Screens in HepG2/C3A cells"

*Supplementary Fig.s:*

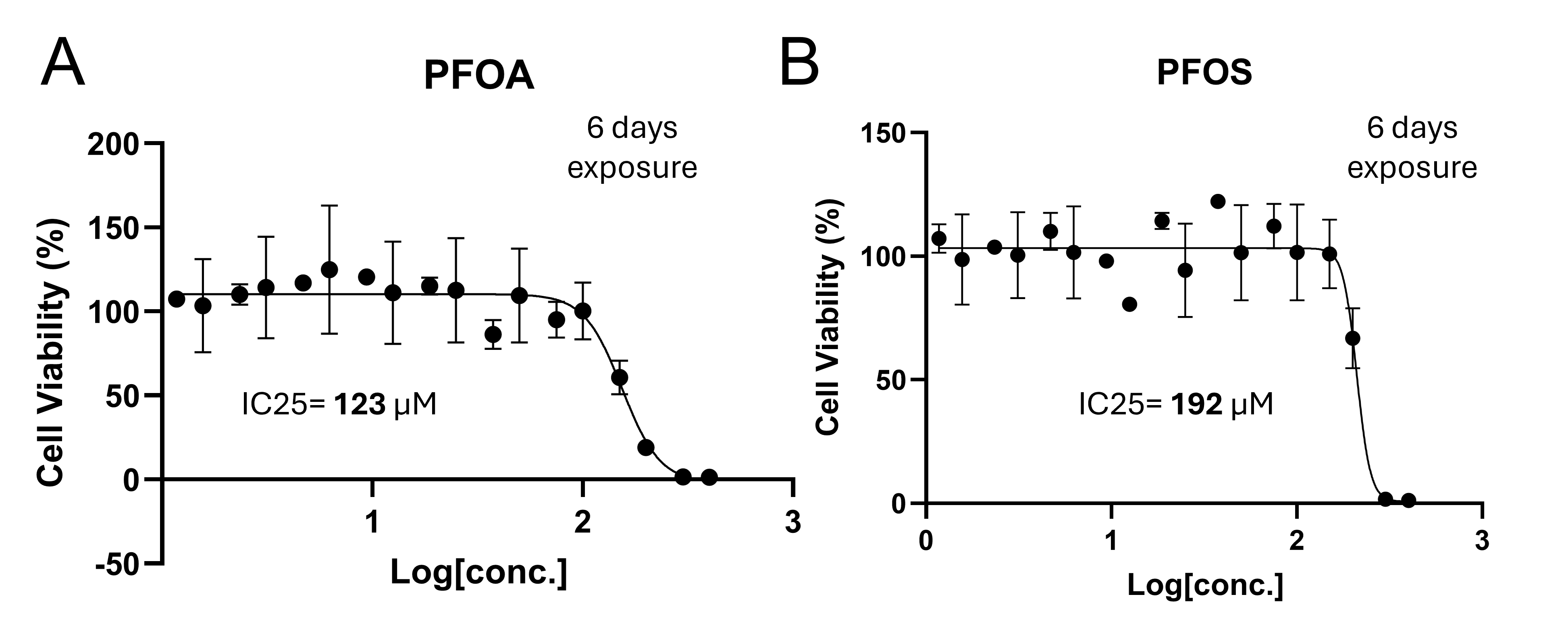

**Fig. S1**. Cell viability of HepG2/C3A cells exposed to PFOS was assessed in a dose-response manner. The IC25 (inhibitory concentration at which the chemical reduces cell viability by 25%) value was calculated using the dose-response sigmoid function in GraphPad Prism. The IC25 of PFOS was 192 µM for 6-day exposure.

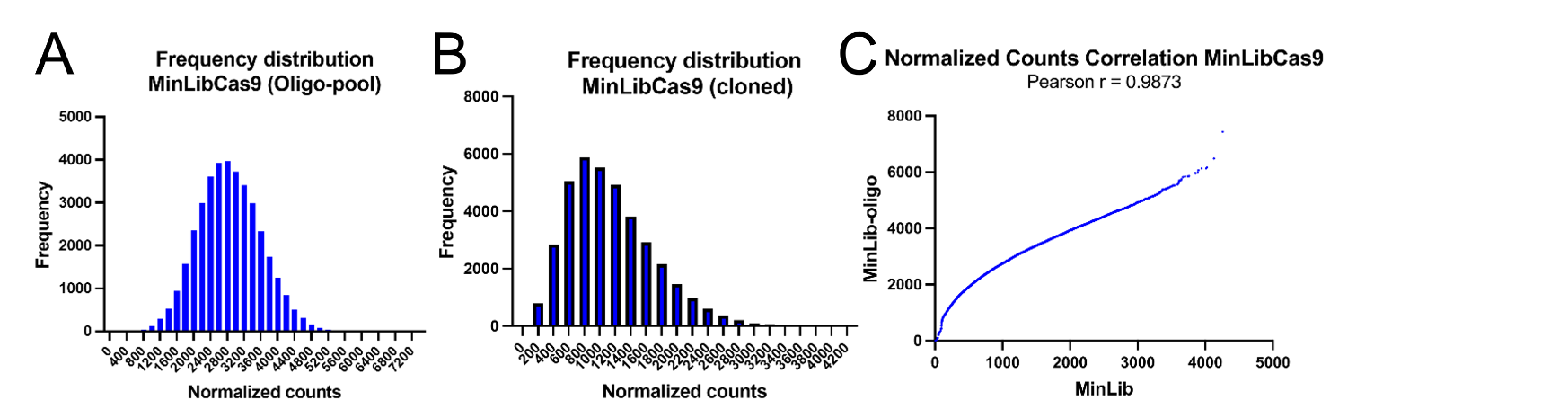

**Fig. S2**. Quality assessment of the custom human minimal genome-wide sgRNA library (MinLibCas9) was conducted. The distribution of normalized sgRNA counts and their corresponding frequencies in the library before and after cloning (Oligo-pool vs. cloned) are shown in panels A and B. The correlation between the Oligo-pool and the cloned library is presented with the Pearson correlation coefficient (r) in panel C.

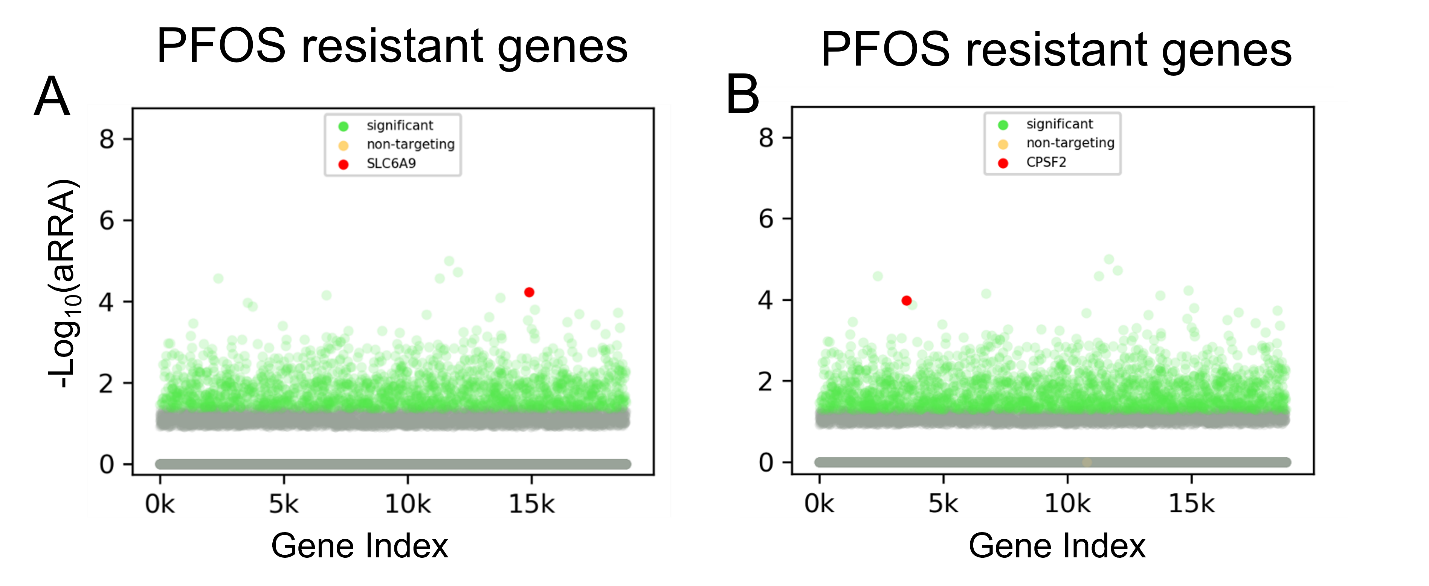

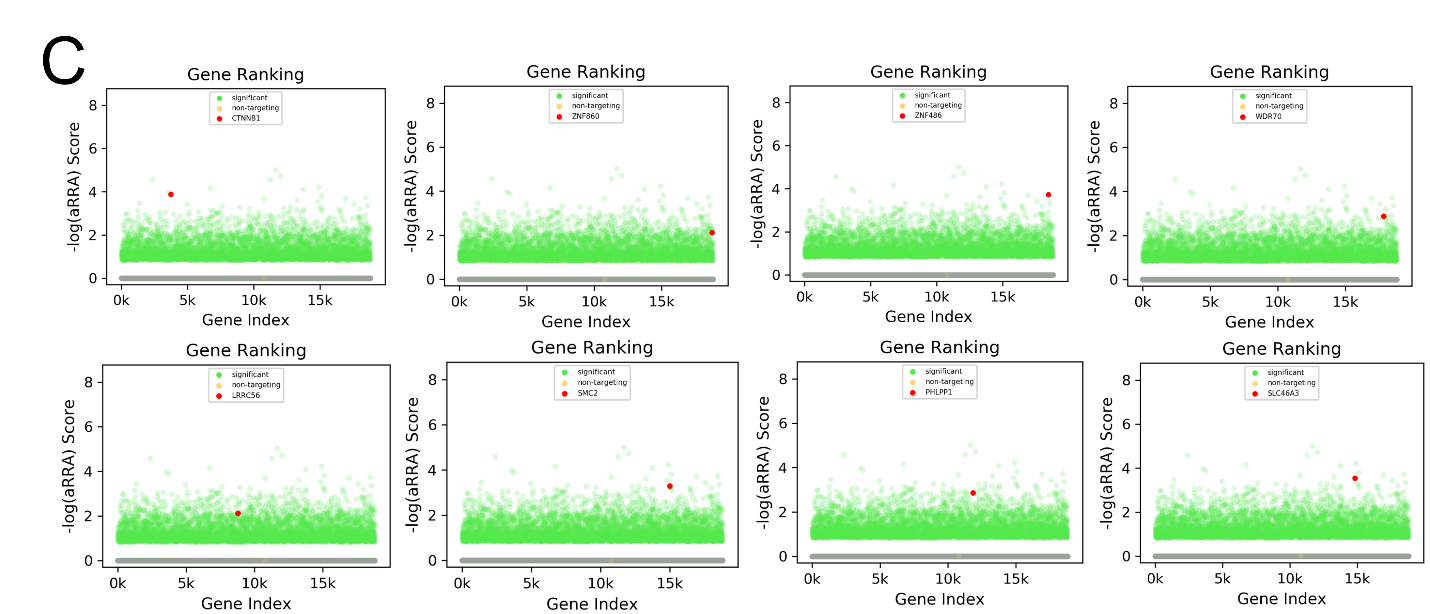

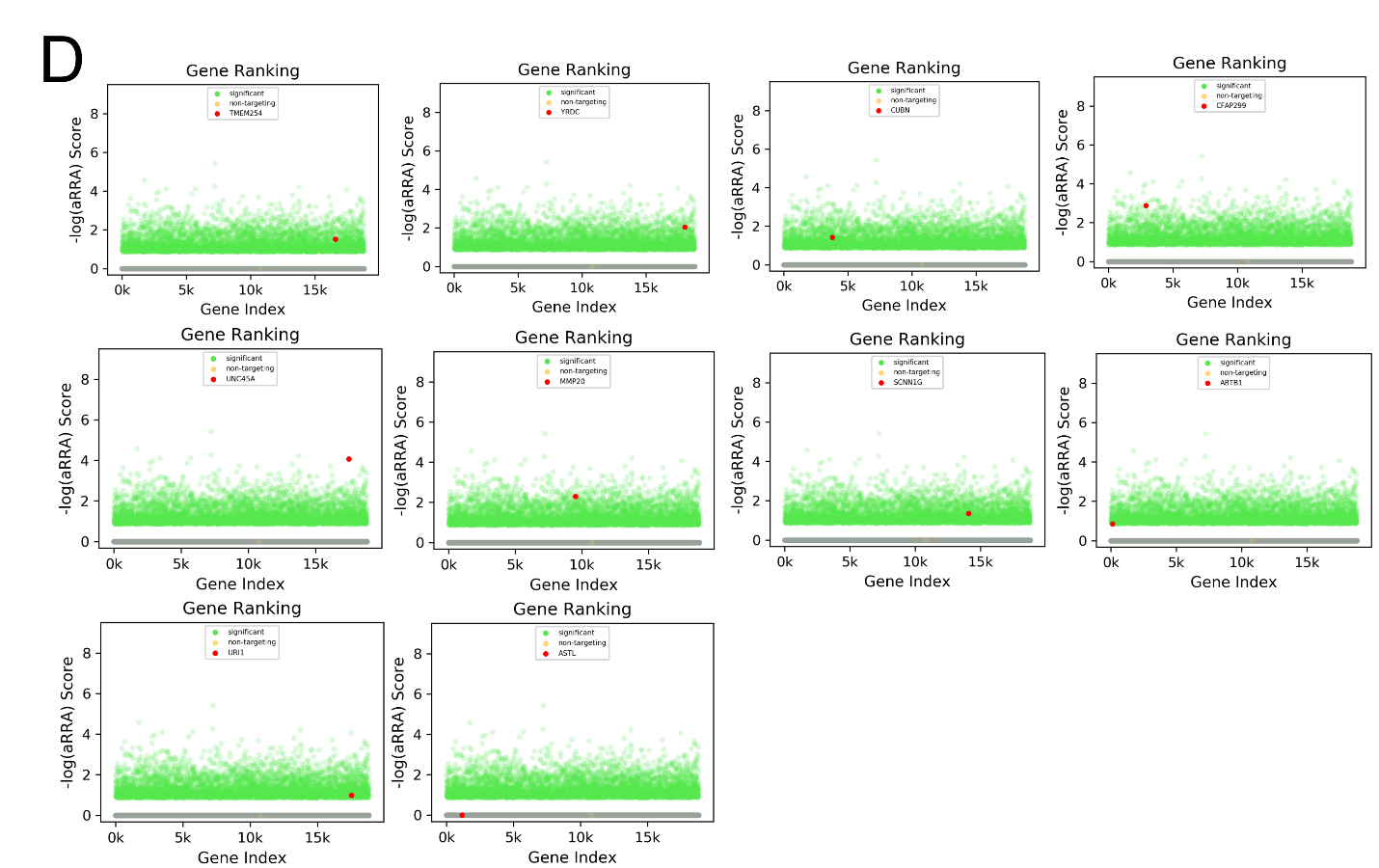

**Fig. S3**. Gene ranking plots from the PinAPL-Py analysis highlight significant positive (resistant) genes responsible for PFOS exposure. The gene ranking plot for PFOS exposure confirms that *SLC6A9* (A) and *CPSF2* (B) were identified as significant genes, both ranked among the top resistant genes. Panel (C) displays the results of the other PFOS resistant top 10 genes while Panel (D) displays PFOS sensitive top 10 genes. The x-axis represents the gene index number, and the y-axis shows the -Log_10_(aRRA) score. aRRA: alpha robust ranking aggregation. Green dots indicate significant genes, grey dots represent non-significant genes, and yellow dots correspond to non-targeting controls, although they may be masked by highly dense grey dots.

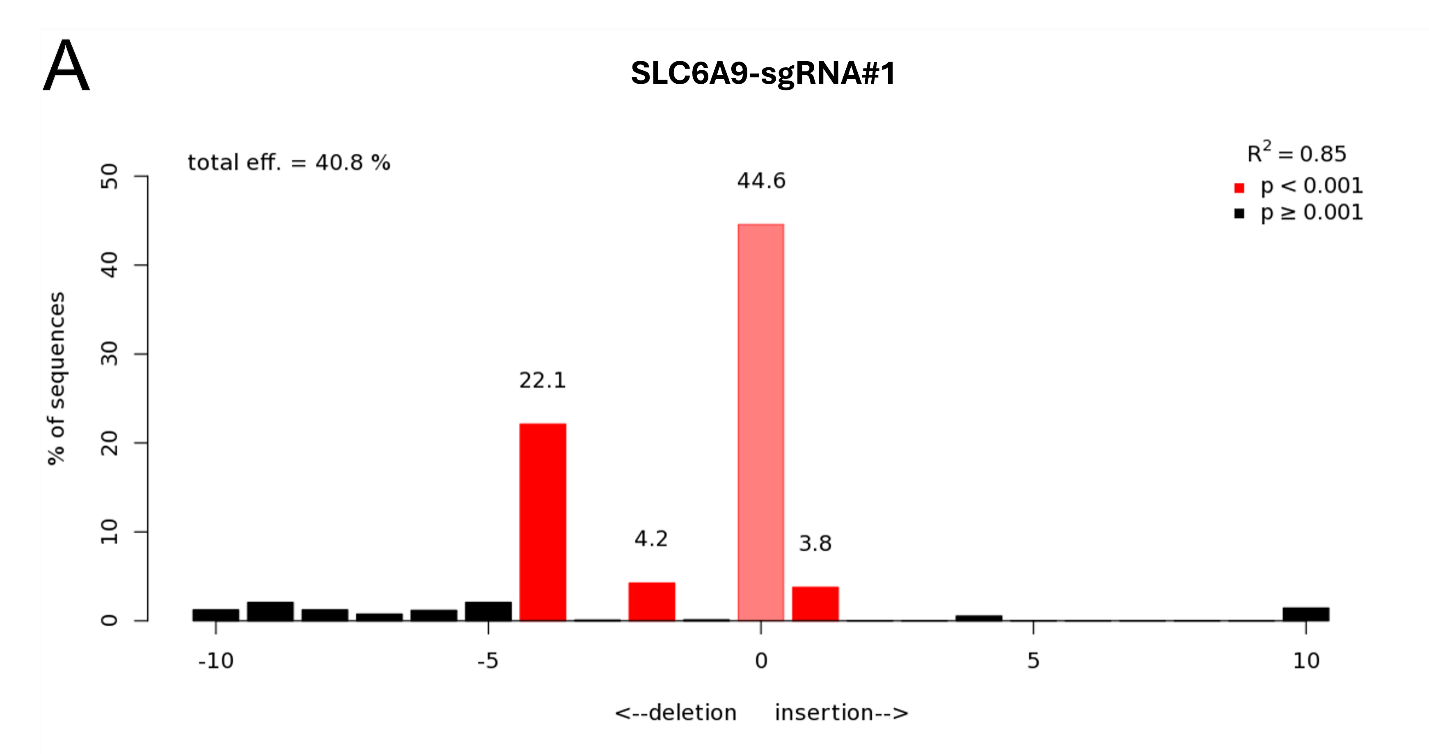

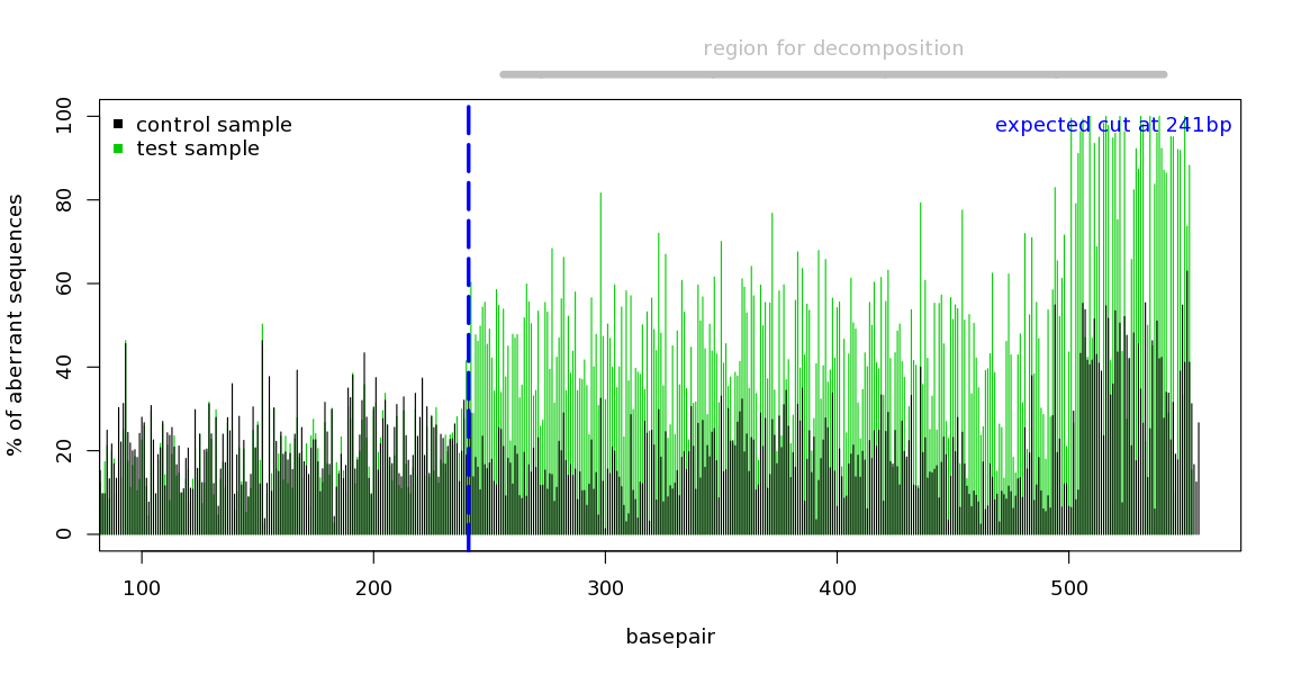

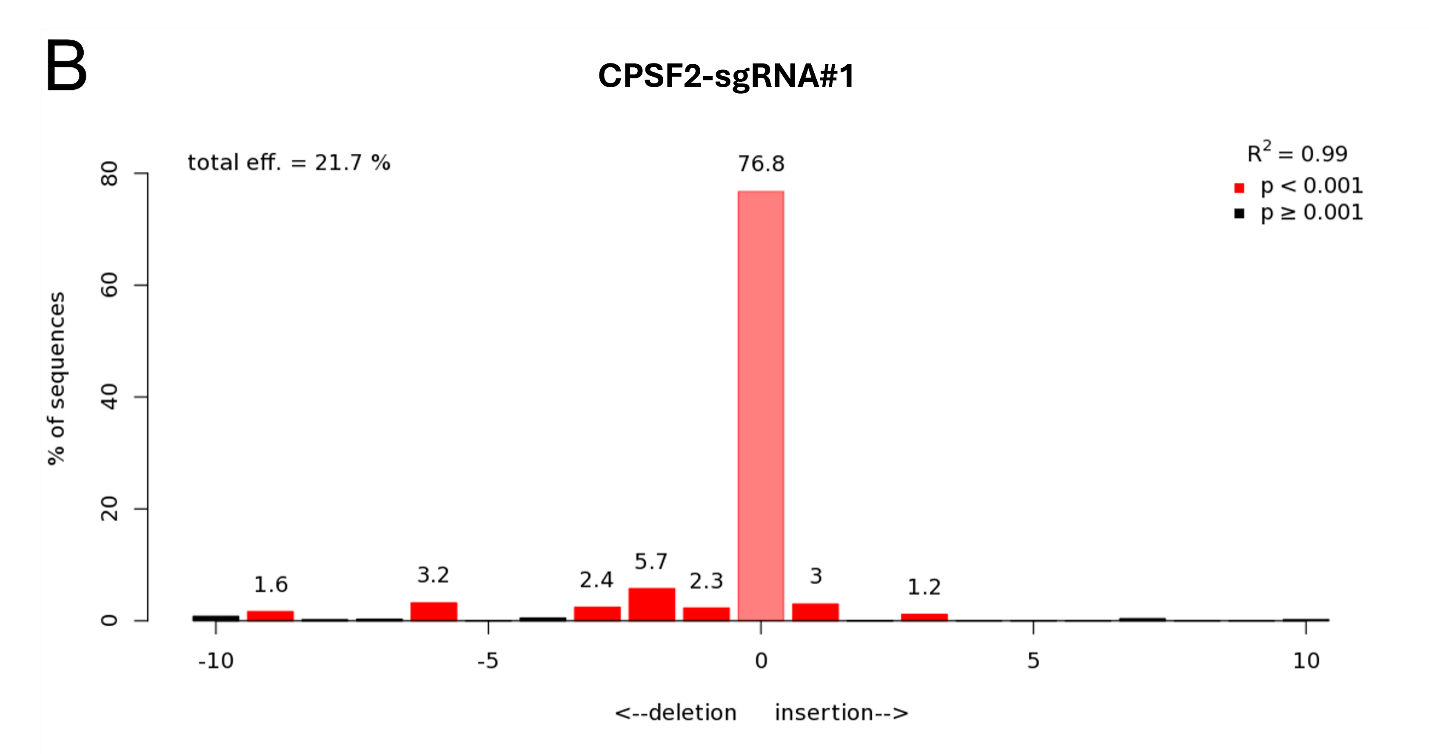

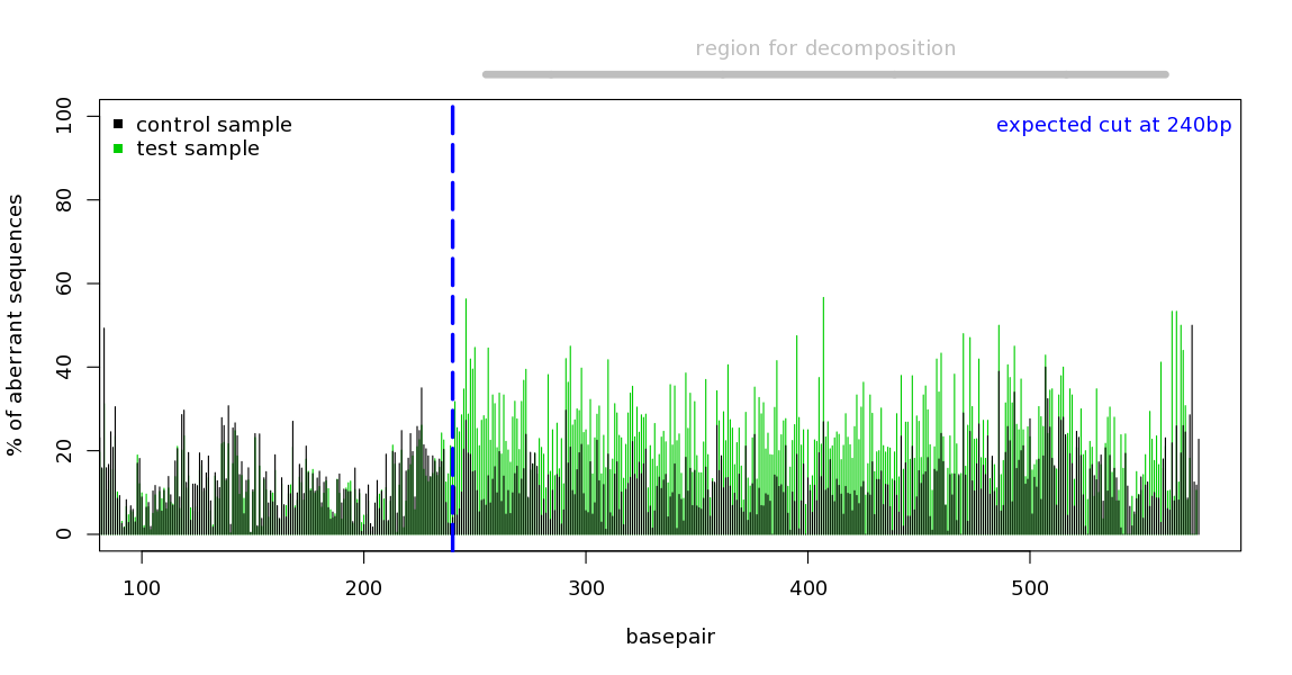

**Fig. S4.** Individual gene knockout results show a percentage of gene-level disruption with TIDE analysis using Sanger sequencing data. (A) *SLC6A9* with sgRNA#1 (total knockout efficiency= 40.8 %), (B) *CPSF2* with sgRNA#1 (total knockout efficiency= 21.7 %).

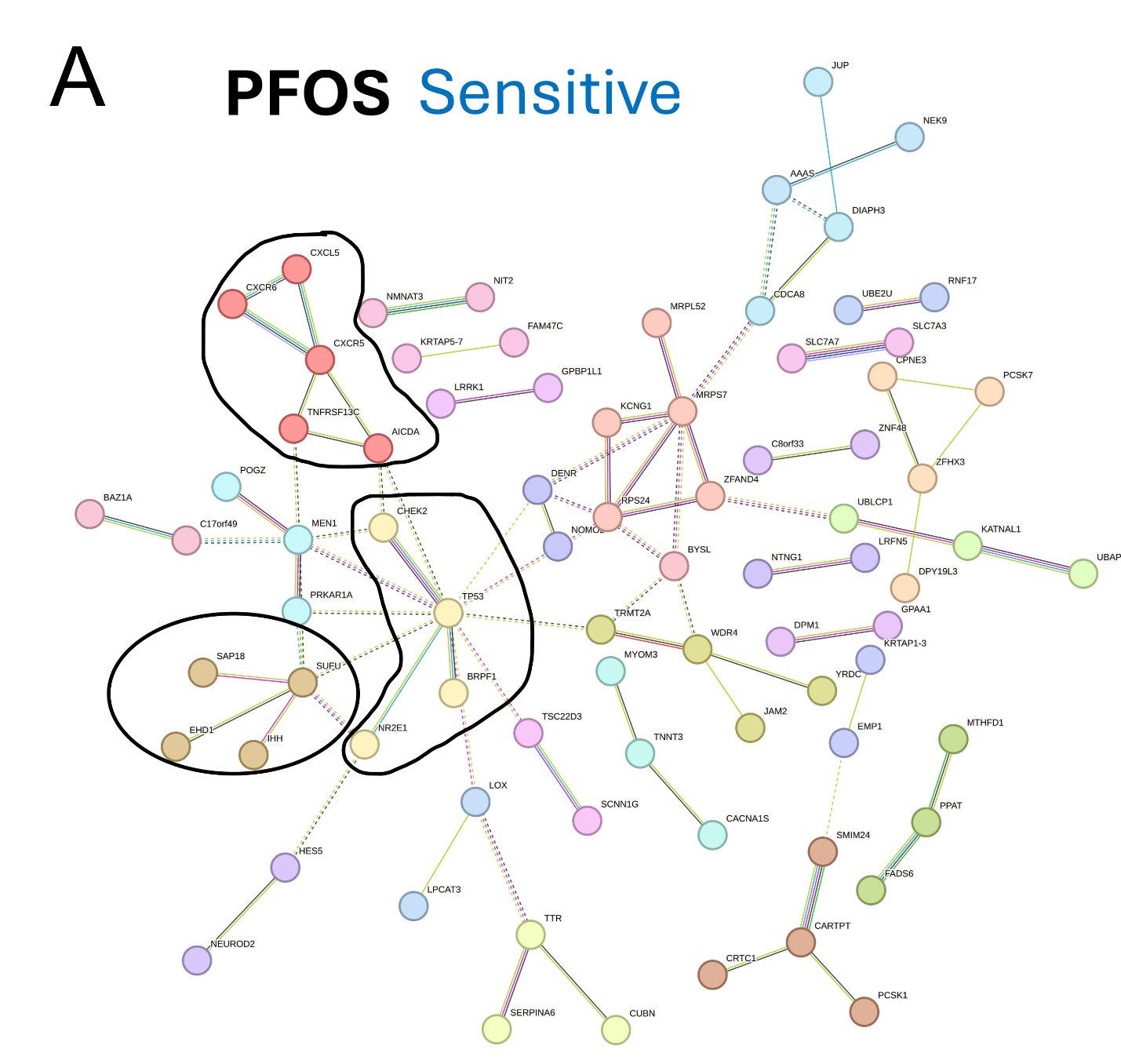

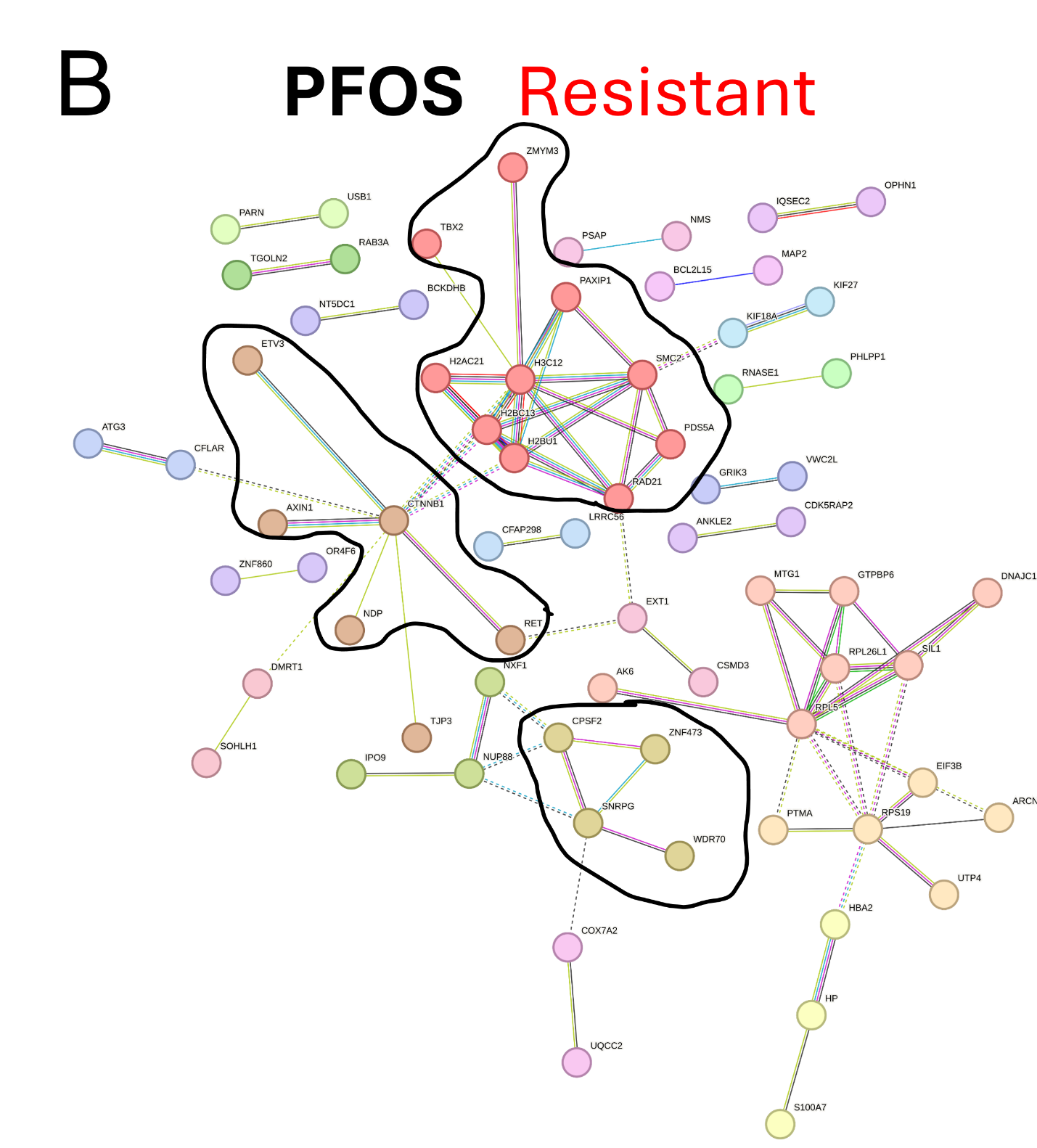

**Fig. S5**. Whole connected STRING clusters of PFOS candidate genes in addition to top three selected clusters (presented in the main Fig.s) for each PFOS candidate gene set. (A) Entire connected clusters of PFOS-sensitive genes in the STRING network. (B) Entire connected clusters of PFOS-resistant genes in the STRING network.

*Supplementary Material and Methods details:*

**HEK29T and HepG2/C3A culture conditions**

The HEK293T cells were cultured in Dulbecco’s Modified Eagle Media (DMEM; Thermo Fisher Scientific, Waltham, MA) and the HepG2/C3A cells were cultured in Minimum Essential Medium (MEM, Gibco). The complete media were supplemented with 10 % fetal bovine serum (FBS, Thermo Fisher Scientific, Waltham, MA), 1X MEM Non-Essential Amino Acids Solution 100X (Gibco, added for HepG2/C3A) and 1X Antibiotic-Antimycotic 100X (Gibco). Cells were cultured in a humidified incubator (Forma™ Series II Water-Jacketed CO2 Incubator, Thermo Scientific™) with 5% CO_2_ at 37 °C.

**PFOS cytotoxicity**

HepG2/C3A cells were seeded at a density of 12,000 cells/well in white-walled 96-well cell culture plates a day prior to exposures. At the day of exposure, attached cells were exposed to a range of nominal concentrations (0–400 mM) of PFOS for 144 h (6 days exposure). To measure the ATP content in wells, 100 µl of the CellTiter-Glo2.0 reagent (the same volume of the culture media) was added to each well containing cells and media and the plate was incubated for 15 min at room temperature in the dark to produce and stabilize the luminescence signal (relative light unit, RLU). The signals were read on a Synergy H1 microplate reader (BioTek Instruments, Winooski, VT). Relative luminescence signals were calculated and subsequently converted to % of viable cells compared to DMSO controls (PFOS concentration: 0 µM). The % values were used to determine 25 % inhibitory concentrations (IC_25_) for PFOS using GraphPad Prism (version 10.1.0) dose-response sigmoid function.

**Preparation and quality check of minimal genome-wide sgRNA library**

We added a few genes that appeared to be missing in the original MinLibCas9 library as well as five different safe harbor control sgRNAs. Of note, we wanted to have the minimal library cloned into a one vector system where both sgRNA and Cas9 are co-expressed to simplify the screening (i.e., bypassing the need to create a Cas9 expressing cell line). Briefly, two previously identified sgRNAs for 18,819 genes from existing sgRNA data (with some annotation corrections) as well as 200 non-targeting controls were synthesized (Twist Biosciences) and the resulting oligo pool was cloned into LentiCRISPRv2_Puro (Addgene #52961) as previously described(Shalem *et al.*, 2014; Sobh *et al.*, 2019). See Supplementary Table S1 for the complete list of sgRNA sequences. The resulting library was verified by next-generation sequencing (NGS) (Illumina NovaSeqX). The sgRNA representation before and after cloning (oligo pool vs. plasmid pool) was verified by NGS of each pool, demonstrating that the distribution of sgRNA counts was similar between the MinLibCas9-oligo (prior to cloning) and MinLibCas9-cloned (post-cloning) samples, with a Pearson correlation coefficient (r) of 0.9873 (Fig. S2). We evaluated the histogram of sgRNA distribution and cumulative sgRNA read counts to assess representation. We consider a six-fold difference between the 90^th^ and 10^th^ percentile as the minimum acceptable distribution of sgRNA representation. The NGS was carried out at the Interdisciplinary Center for Biotechnology Research (ICBR), University of Florida at Gainesville, using the NovaSeqX paired 150 bp high-throughput platform (Illumina).

**Lentiviral production and transduction**

HEK293T cells were used to produce lentivirus by co-transfection of the MibLibCas9 library plasmids, the packaging plasmid psPAX2 (#12260, Addgene), and envelope plasmid pMD2.G (#12259, Addgene). The MinLibCas9 lentiviral library was functionally tittered in HepG2/C3A cells to determine the amount of virus required to obtain a multiplicity of infection of 0.3-0.5. For large-scale transduction for the screens, HepG2/C3A cells were seeded in four 12-well culture plates (1x106 cells/well). After a 24 h incubation period, polybrene (Sigma) was added at a concentration of 8 µg/ml along with 0.5 µl (MOI= 0.3) of the titrated MinLibCas9 lentiviral library in each well, followed by centrifugation (spinfection) at 33 ◦C at 1000xG for 2 h. After further incubation at 37 ◦C for 30 h, the lentiviral mix was replaced with 1 mL of the growth media in each well. After 48 h recovery period, the pooled cells were treated with puromycin (2 μg/mL) for 5 days to enrich for transduced cells before chemical exposures, resulting in almost no cells in the non-transduced control flask.

**Single gene knockout validations of candidate genes of PFOS exposure**

To produce lentiviruses, LentiCRISPRv2 constructs harboring each of the *SLC6A9* and *CPSF2* sgRNAs were co-transfected in HEK293T cells with the envelope pMD2.G and packaging psPAX2 plasmids. In brief, HEK293T cells were seeded in T25 tissue culture flasks the day prior to co-transfection. One hour before the transfection, the medium was replaced with 2.25 mL of Opti-MEM medium (Gibco). For each of the gRNAs, the DNA mixtures consisted of 3.4 μg lentiCRISPRv2-gene target plasmid, 1.7 μg pMD2.G, and 2.6 μg psPAX2 diluted in 700 μL Opti-MEM and 35 μL plus reagent (Thermo Scientific), and the lipofectamine mix consisted of 700 μL Opti-MEM and 17.25 μL lipofectamine 2000 (Thermo Scientific). The two mixtures were incubated for 5 min at room temperature and then combined and incubated at room temperature for 20 min. The lipofectamine-DNA complex was added to the HEK293T cells, and the cells were then incubated for 6 h at 37 °C and 5% CO_2_. The transfection medium was replaced with 6 mL of lentivirus harvest medium (DMEM high glucose, 20% FBS, 1× penicillin/streptomycin). The lentiviruses were harvested 48H post-transfection as follows: cell culture medium was collected and centrifuged at 3000 rpm for 10 min at 4 °C. To concentrate the lentiviruses, 2 mL (1/4 of the supernatant volume) of Lenti-X concentrator (Takara Bio) was mixed with the supernatant. The mixture was incubated on ice for 2 h 30 min and then centrifuged at 1500xG for 45 min at 4 °C. The obtained pellet was resuspended in 300 μL of harvest media and stored at −80 °C until the use. HepG2/C3A cells were transduced separately with each of the SLC6A9 and CPSF2 lentiviral preparations using the spinoculation method. In brief, 2.5 million HepG2/C3A cells were seeded in a 12-well plate in 2 mL of the complete MEM medium. Polybrene (8 μg/mL final concentration) and 10 μL of the lentiviral solution were added to the cells and mixed gently up and down. The 12-well plate was centrifuged at 1000g for 90 min at 34 °C. After centrifugation, the cells were transferred to 15 mL conical tubes and pelleted (300xG for 5 min). The cell pellet was resuspended in 6 mL of the complete MEM medium and cultivated in a T25 tissue culture flask for 48 h. The transduced cells that integrated the lentiviral construct were then selected in the complete MEM medium supplemented with 2 μg/mL puromycin for 6 days.
